# Polymer physics and machine learning reveal a combinatorial code linking chromatin 3D architecture to 1D epigenetics

**DOI:** 10.1101/2021.03.01.433416

**Authors:** Andrea Esposito, Simona Bianco, Andrea M. Chiariello, Alex Abraham, Luca Fiorillo, Mattia Conte, Raffaele Campanile, Mario Nicodemi

## Abstract

The mammalian genome has a complex 3D organization, serving vital functional purposes, yet it remains largely unknown how the multitude of specific DNA contacts, e.g., between transcribed and regulatory regions, is orchestrated by chromatin organizers, such as Transcription Factors. Here, we implement a method combining machine learning and polymer physics to infer from only Hi-C data the genomic 1D arrangement of the minimal set of binding sites sufficient to recapitulate, through only physics, 3D contact patterns genome-wide in human and mouse cells. The inferred binding sites are validated by their predictions on how chromatin refolds in a set of duplications at the *Sox9* locus against available independent cHi-C data, showing that their different phenotypes originate from distinct enhancer hijackings in their 3D structure. Albeit derived from only Hi-C, our binding sites fall in epigenetic classes that well match chromatin states from epigenetic segmentation studies, such as active, poised and repressed states. However, the inferred binding domains have an overlapping, combinatorial organization along chromosomes, missing in epigenetic segmentations, which is required to explain Hi-C contact specificity with high accuracy. In a reverse approach, the epigenetic profile of binding domains provides a code to derive from only epigenetic marks the DNA binding sites and, hence, the 3D architecture, as validated by successful predictions of Hi-C matrices in an independent set of chromosomes. Overall, our results shed light on how complex 3D architectural information is encrypted in 1D epigenetics via the related, combinatorial arrangement of specific binding sites along the genome.

## INTRODUCTION

The genome of higher organisms has a complex spatial organization within the cell nucleus^1–6^ as revealed by recent technologies^7–13^. Chromosomes are folded in a sequence of 0.5-1.0Mb wide domains, named TADs^14,15^, in sub-TADs and loops, and in larger structures such as A/B compartments^7^ and meta-TADs^16^. Importantly, such an organization serves vital functional purposes, as for instance distal enhancers control their target genes by establishing physical contacts with them, disruptions being linked to human diseases^17–19^. However, how chromatin architecture is shaped and orchestrated remains mostly unknown.

To rationalize the complexity of Hi-C data, polymer models from statistical physics^20–32^ and a variety of computational methods^33–36^ have been developed. A class of models, such as the *Strings and Binders* (SBS) model^21^, has focused on the classical scenario where loops and contacts between distal DNA sites are established by diffusing molecules such as Transcription Factors (TFs), or some effective interaction potential, bridging cognate binding sites by thermodynamics mechanisms of phase separation^21,25–32,37–39^. Another interesting classical scenario has been considered by off- equilibrium polymer models where loops are formed by extrusion, e.g., by molecules that bind to DNA and extrude a loop^22–24^, based on prior knowledge of the involved molecular factors, such as CTCF binding sites^29,40^.

Here, we use a machine learning approach (PRISMR^41^) to infer from only Hi-C data the genomic location of the minimal set of binding sites best explaining contact patterns across chromosomes by only polymer physics via the molecular mechanisms envisaged by the SBS model. While PRISMR was previously applied to Mb wide genomic regions, we optimize its performance to extend the approach to the entire genome, improving the statistical power of our method of three orders of magnitude to understand how complex 3D architectural information is encrypted in 1D epigenetic signals. Without prior knowledge of binding factors, our approach can infer genome-wide the specific location of the distinct binding sites whereby DNA contacts are established as captured by Hi-C data, returning a picture of the key elements underlying chromosomes folding. We show that the SBS polymer model informed with the inferred binding sites recapitulates Hi-C data across chromosomes in human^42^ and murine^14^ cells with high accuracy, illustrating that its minimal ingredients are sufficient to make sense of a substantial fraction of contact patterns genome-wide. For sake of simplicity, we focus on the SBS model, but the method can be extended to accommodate additional mechanisms, such as loop extrusion^22–24^.

To test the inferred binding sites of the model and its envisaged folding mechanisms, we compared its predictions about the impact of mutations on chromosome conformations against independent experimental data. As a case study, we considered the *Sox9* locus, where cHi-C data are available for a set of duplications^43^. We implemented in the wild-type chromosome model those duplications and derived *de novo* the corresponding contact maps that are successfully compared to cHi-C data, with no fitting parameters available. Our analysis also shows that different genomic variants produce different neo-TADs around *Sox9* marked by specific enhancer hijackings, hence resulting in different phenotypes.

Importantly, our inference procedure does not exploit previous knowledge on binding factors or epigenetics marks. Hence, the inferred binding domains can be used to bring together independently derived information on architecture and epigenetics, e.g., by crossing their genomic position with ENCODE databases. We find that the different binding domains fall in similarity classes based on epigenetics, well matching functional chromatin states derived in linear epigenetic segmentation studies such as active, poised and repressed states^10,44–47^. However, we discover that they have an overlapping, combinatorial genomic distribution at the current resolution of Hi-C experiments, lacking in linear segmentation studies, which is shown to be required to explain Hi-C contacts with high accuracy genome-wide.

Finally, we validated the discovered association between machine learned binding sites and epigenetic features by reversing the approach. In the considered cell types, we used the epigenetic profiles of the different binding domains of even chromosomes as a code to derive from only histone marks of odd chromosomes the location and type of their binding sites. Next, those binding sites were used to inform the polymer models of odd chromosomes, which predicted the corresponding Hi-C matrices with an accuracy comparable to those directly inferred from Hi-C data.

Overall, our results provide insights on how the 1D combinatorial arrangement of a comparatively small number of binding site types, barcoded by distinctive epigenetic signatures, encodes the architectural information guiding chromatin organizing factors to establish, through physics, the multitude of specific 3D contacts across chromosomal scales.

## RESULTS

### Inferred binding domains explain Hi-C data genome-wide

To dissect the molecular mechanisms that contribute to chromatin folding, we used the PRISMR machine learning procedure^41^ to infer the minimal SBS polymer model best explaining *in situ* Hi-C contact maps in human GM12878 B-lymphoblastoid cells at 5kb resolution across chromosomes^42^(**Fig. 1B, Fig. S1**, Materials and Methods). In the *String and Binders* (SBS) model^21^, a chromatin filament is modeled as a self-avoiding string of beads, including specific binding sites for diffusing molecules; such binders can bridge distal cognate sites along the sequence, producing loops and physical contacts (**Fig. 1C**). In particular, in the SBS model, contact domains of homologous sites are spontaneously established by their cognate binders via a thermodynamic mechanism known as micro-phase separation^26,30,39^. Specifically, PRISMR finds the minimal combination of the binding sites of the SBS model (**Fig. 1D**) that reproduces, within a given accuracy threshold, the experimental Hi-C contact matrix based only on polymer thermodynamics (Materials and Methods). The different groups of homologous binding sites are named the *binding domains* of the model. Importantly, PRISMR uses just Hi-C data as input, with no prior knowledge of binding factors.

**Fig.1.**
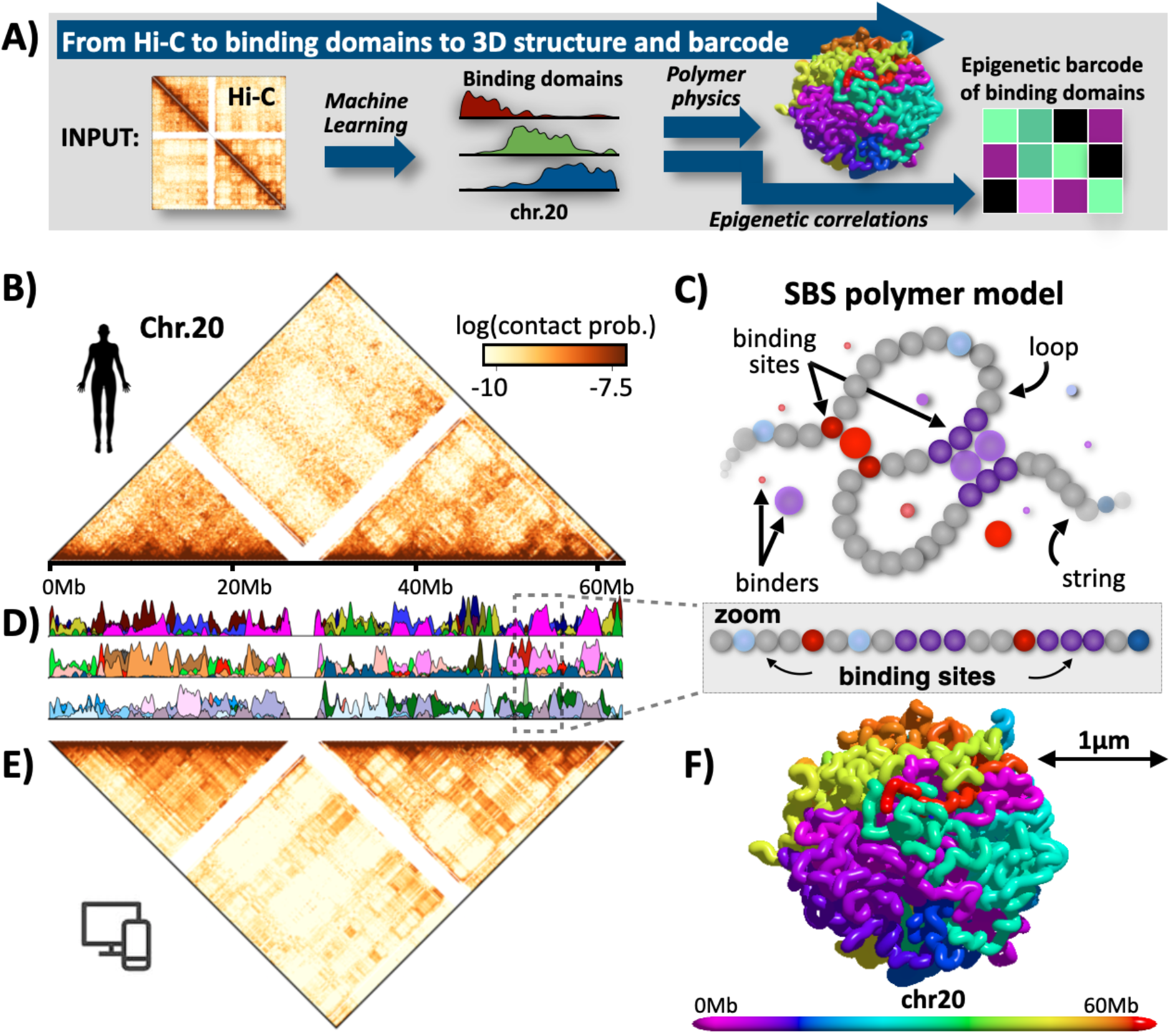
The inferred binding domains explain Hi-C data across chromosomes. **(A)** Our method combines machine learning and polymer physics to infer from only Hi-C data the genomic location of the minimal set of binding sites required to recapitulate chromatin conformations genome-wide by use of the SBS polymer model of chromatin. Additionally, by correlations with epigenetic data, the inferred binding domains can be assigned a molecular barcode. **(B)** *In situ* Hi-C data^42^ of chromosome 20 at 5kb resolution (log-scale). **(C)** A scheme of the SBS polymer model of chromatin: it quantifies the scenario where diffusing binders bridge and loop distal cognate binding sites. Each colored bead is a single binding site. The genomic location of the binding sites encodes the 1D information whereby their cognate binders produce the 3D structure via polymer physics. **(D)** Plots displaying the position and abundance of the different types of binding sites (binding domains) along chromosome 20, as inferred by our method. For visualization purposes, the different domains, each represented with a different color, are drawn in groups of 10 in different rows. Albeit derived from only Hi-C data, the binding domains have specific correlations each with a set of epigenetic marks, and the colors reflect those associations (see Fig.3). **(E)** The model inferred contact matrix of chromosome 20 has a Pearson, distance-corrected Pearson and stratum adjusted correlation with Hi-C respectively equal to r=0.97, r’=0.85, SCC=0.92. Similar results are found across chromosomes (Fig. S1). **(F)** A time snapshot of the 3D structure of the SBS model of chromosome 20.

To check whether the model can explain *in situ* Hi-C data genome-wide, we compared the PRISMR derived SBS contact matrices to the original data (**Fig. 1E, Fig. S1**). In particular, we computed their Pearson correlation coefficient, r, their distance corrected Pearson correlation coefficient, r’, and their HiCRep stratum adjusted correlation coefficient, SCC^48^. The last two measures were considered to account for genomic proximity effects (see Materials and Methods). Model and experimental data were found to be comparatively similar across chromosomes, as r, r’ and SCC range around r=0.94, r’=0.74 and SCC=0.86, respectively. Notably, from the SBS model the thermodynamics ensemble of chromosomal 3D conformations can also be derived; a snapshot, e.g., of chromosome 20 is pictured in **Fig. 1F**.

Additionally, to prove the general validity of the method, we tested its performance on a mouse embryonic stem cells Hi-C dataset at 40kb resolution^14^, finding that the PRISMR inferred and experimental contact matrices have high correlation values, comparable to those reported above for the 5kb human data (**Fig. S2**).

The model binding domains, i.e., the sets of homologous binding sites along the chromosomes, are the output of PRISMR. The algorithm returns 30 different binding site types per chromosome in GM12878 (**Fig. 1D**, see Materials and Methods). Interestingly, a single binding domain covers on average a genomic length comparable to the size of a TAD (3.1+-1.9Mb), yet the range of their interactions, *r*_*Int*_, extends more than one order of magnitude longer, up to tens of Mbs (**Fig. S3A**, Materials and Methods). The distribution of *r*_*Int*_, *P(r*_*Int*_), is significantly different from a random control model obtained by bootstrapping the location of binding sites (Materials and Methods) and is asymptotically consistent with a power-law scaling, *P ∼ 1/r*_*Int*_, typical of hierarchical structures made of domains within domains, as in Cantor sets (**Fig. S3A**, Materials and Methods). The broad range of values of *r*^*Int*^ shows that chromatin interactions extend above the size of single TADs, with higher-order 3D structures formed at scales below and above the A/B compartment level^16^. The derived 3D structure of chromosomes (**Fig. 1F**) shows indeed that, rather than being a linear chain of TADs, they tend to fold on themselves in complex structures, such as meta-TADs^16^ (see also ^31^).

Taken together, the high correlations found between the SBS model and Hi-C contact data show that the 1D binding domains inferred genome-wide by PRISMR contain key information sufficient to recapitulate 3D contact patterns genome-wide in human and mouse cells. That sheds light on the molecular mechanisms shaping chromosome architecture, supporting the view that the combinatorial action of a comparatively small number of TFs, mediating the interactions between cognate binding sites, can spontaneously fold chromatin in its 3D structure through just the laws of physics.

### Validation of the inferred binding domains against duplications in the *Sox9* locus

To validate the binding domains inferred by our approach, i.e., the determinants of folding and their envisaged mode of action, we compared our model predictions against previous independently produced cHi-C data in E12.5 limb bud cells from mice carrying homozygous structural variants in the *Sox9* locus^43^. We considered three mutations (**Fig. 2, Fig. S4**): a 0.4Mb duplication (*DupS*) in the non-coding DNA region within the *Sox9* gene TAD (intra-TAD duplication), associated with female to male sex reversal in humans; a 1.6Mb duplication (*DupL*) that encompasses the neighboring TAD boundary, with no phenotypic effect; and a slightly longer 1.8Mb duplication (*DupC*), associated to limb malformation, which also includes *Kcnj2*, the next flanking gene. Specifically, we implemented those mutations in the SBS polymer model of the wild-type region in mESC inferred by PRISMR^30^ and derived the novel contact matrices from only polymer physics, with no fitting parameters whatsoever. The Pearson and distance-corrected Pearson correlation coefficients between the model predicted and cHi-C contact maps across the three mutations are as high as r=0.88 and r’=0.48, reflecting their good degree of similarity (**Fig. 2, Fig. S4**).

**Fig.2.**
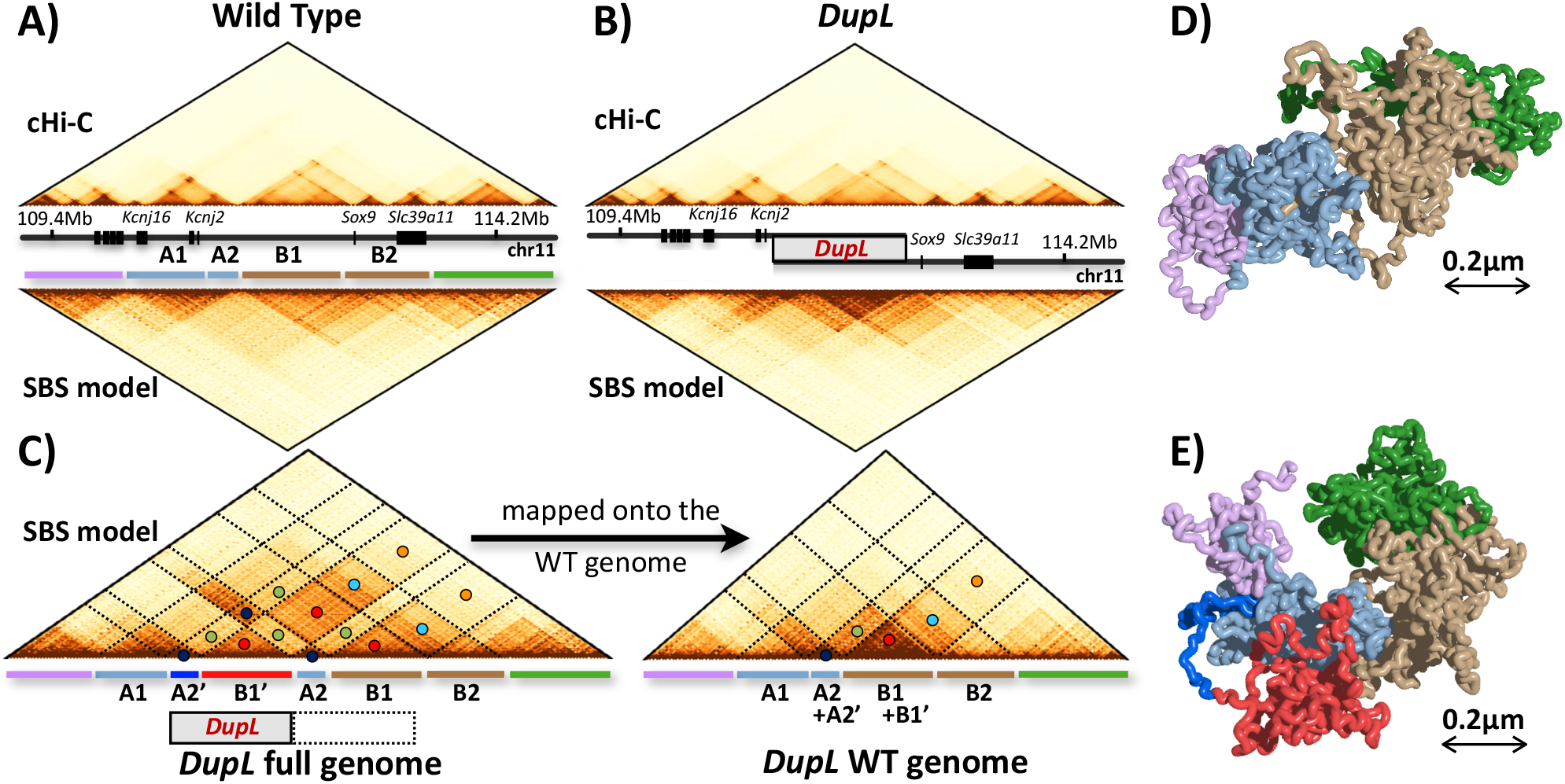
The inferred binding domains are validated against mutations at the Sox9 locus. **(A)** Contact data^43^ of the wild type *Sox9* locus from cHi-C experiments in E12.5 limb buds (top) and of the SBS model of the locus in mESC (bottom) have a correlation r=0.89 and r’=0.44. **(B)** Based on the WT model, the contact map of a mutant bearing the *DupL* duplication is predicted from only physics (bottom). It has a good correlation (r=0.82, r’=0.41) with independent *DupL* cHi-C data^43^ (top). Model predictions are also validated across the other available *Sox9* mutations (Fig. S3, S4). **(C)** Mapping the model contacts on the *DupL* full genome clarifies the origin of the associated neo-TAD (red). The colored circles mark corresponding interaction regions as mapped on the WT and *DupL* full genomes. **(D)-(E)** Snapshots of the model predicted 3D conformation of respectively the WT and *DupL* locus (the color scheme reflects the colored bars in panel A and C) with its neo-TAD. Different mutations result in different 3D structures and distinct enhancer-hijackings, explaining their phenotypes (Fig. S3, S4).

**Fig.3.**
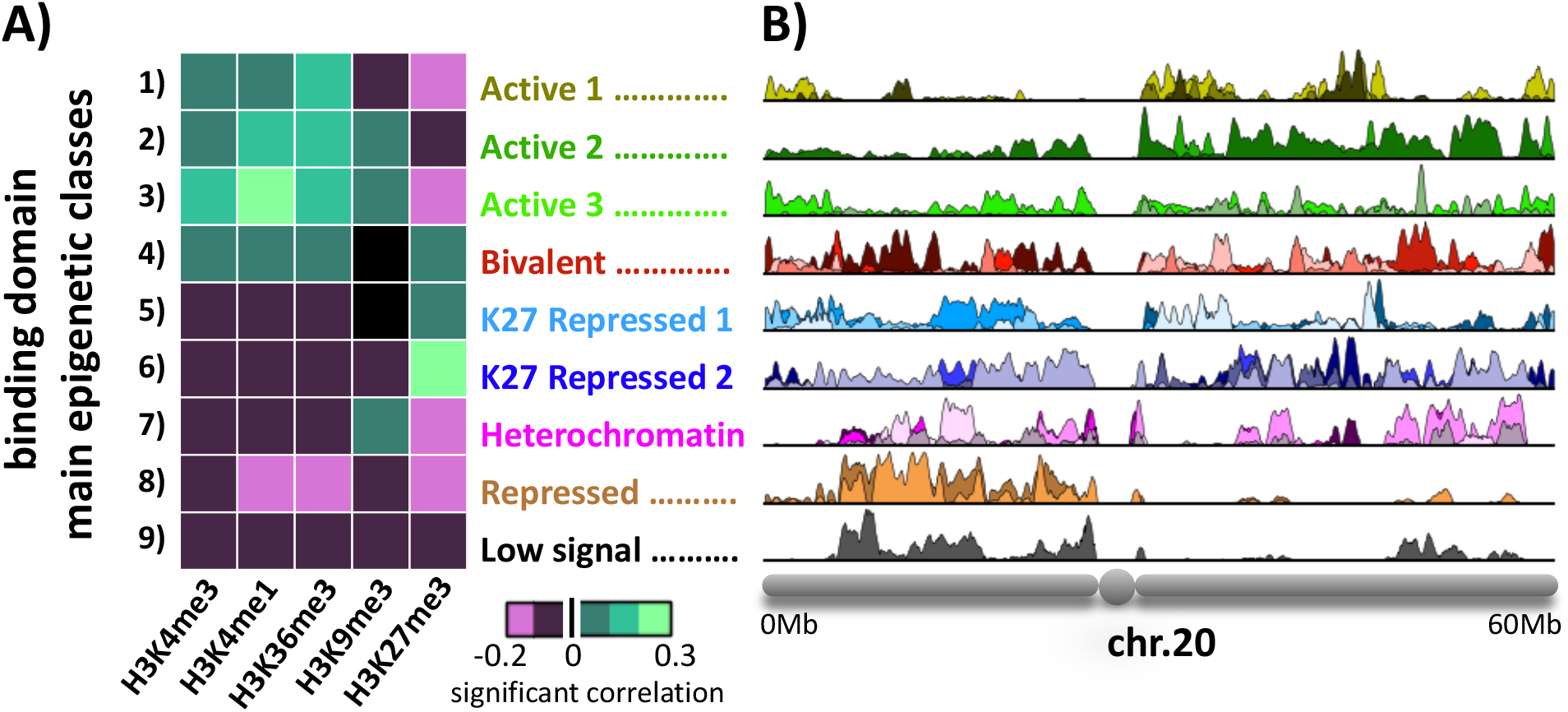
Epigenetic profiles of the inferred binding domains. **(A)** The model binding domains, inferred from Hi-C data only, correlate each with a specific set of epigenetic tracks. They cluster in 9 main classes genome-wide according to their correlations with the shown ENCODE key histone marks (Fig. S6). The epigenetic profile, i.e., the barcode of the centroid of each class is shown in the heat-map. The 9 classes match well chromatin states derived in epigenetic segmentation studies. **(B)** Their abundance along chromosomes is not uniform (p-value<0.05, Fig.7), as shown here for the binding sites of chromosome 20.

Those results provide a stringent validation to our approach and demonstrate that predictions on the 3D structure of chromatin based on the inferred binding domains can be accurate to the point to anticipate ectopic contacts produced by disease-associated mutations.

### Enhancer hijackings in neo-TADs at the mutated *Sox9* locus link to phenotype

To understand the origin of the ectopic contacts in the mutated systems, within our model we dissected the interactions of the duplicated from the original DNA sequences and the corresponding 3D structures, pieces of information inaccessible by Hi-C data (**Fig. 2, Fig. S4, S5**).

*DupS* is fully included within the TAD encompassing *Sox9* (**Fig. S4A**). Within our model, a TAD and its corresponding enrichment of interactions derive from the presence of a prevailing type of binding sites in that DNA region (see, e.g., TAD *A, B, C* in **Fig. 4D**-**F**). Hence, the duplicated and the original sequence in *DupS* (region B2’ and B2 in **Fig. S5A**) share many homologous binding sites, which produce the contacts between such regions visible in the interaction matrix mapped along the full, duplicated genome (**Fig. S4A, S5A**). When those contacts are mapped back onto the wild-type sequence, an excess of interactions appears localized around the mutated region within the corresponding TAD, but no major changes to the overall contact pattern, as experimentally found in cHi-C data^43^. The model derived 3D structure of the mutated locus shows, indeed, that the duplicated region remains well embedded into the rest of the locus (**Fig. S5C**).

**Fig.4.**
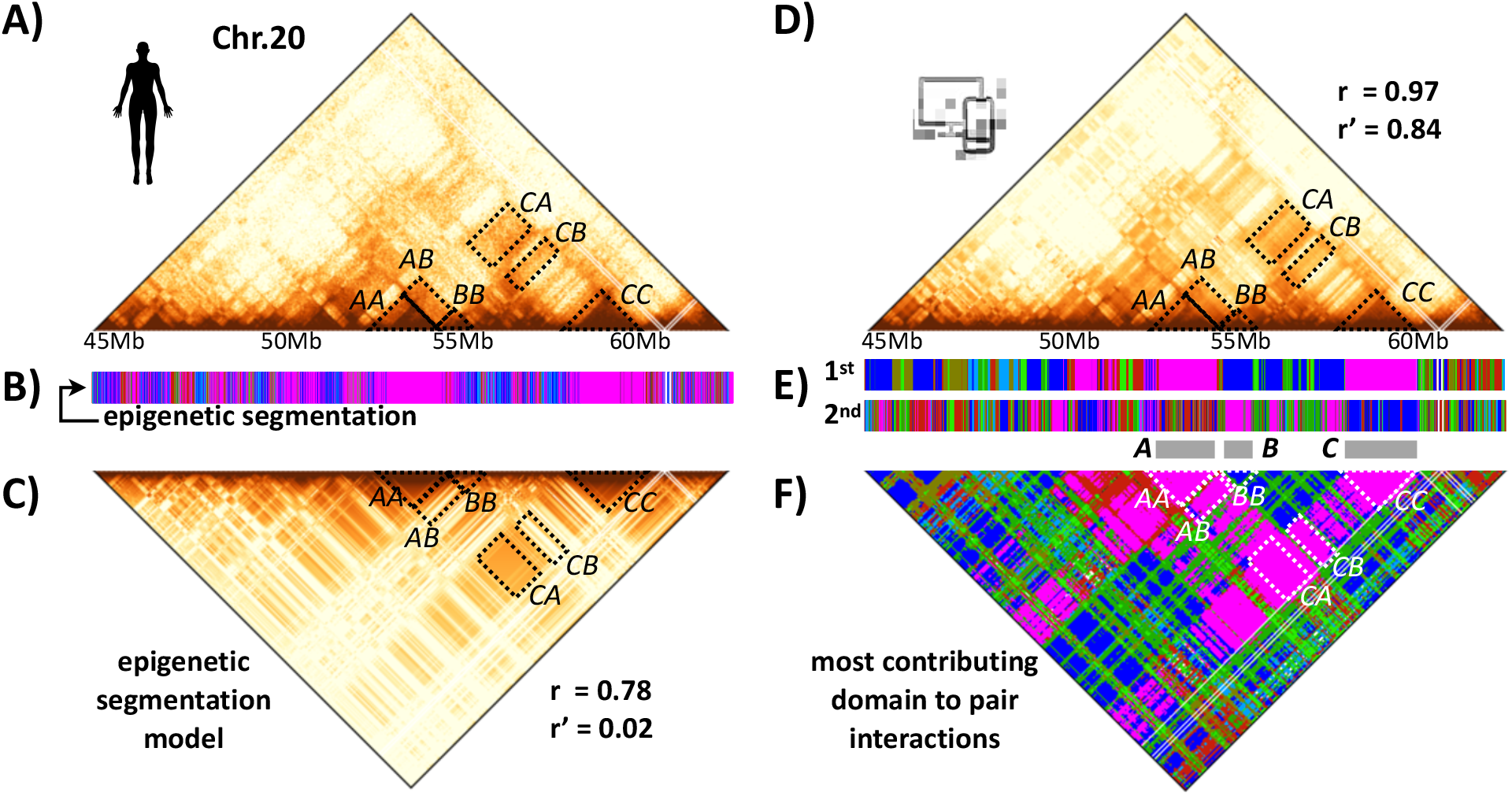
Chromatin architecture patterns are only partially captured by linear epigenetic segmentation. *In situ* Hi-C data^42^ (scales as in Fig. 1) of a 20Mb wide region on chr20 in GM12878 and its linear epigenetic segmentation are shown. **(C)** The contact map of a model based only on homotypic interactions between linear segmented epigenetic domains has a Pearson correlation r=0.78 with Hi-C data. Yet, its distance corrected correlation is much lower, r’=0.02, returning only a marginal improvement over a control model where each interaction is replaced by the average at the corresponding genomic separation. **(D)** The contact map of the inferred SBS model of the region has r=0.97 and r’=0.84 with Hi-C data. **(E)** The PRISMR inferred 1st and 2nd most abundant binding site types of the SBS model of that region are shown. **(F)** The plot of the SBS most contributing binding domain to each pairwise contact highlights that a combinatorial overlap of different binding site types along the sequence, missing in linear segmentations, is required to capture the complexity and specificity of interaction patterns. For example, interactions (CC) within the TAD in region C are mainly related to binding domains in class 7 (magenta), the most abundant one in *C. A* and *C* also interact mainly through class 7, the most abundant in *A* too. Yet, region *B*, where class 6 (dark blue) is the most abundant, interacts with *C* mainly through class 7, its 2nd most abundant. Analogously, contacts between *A* and *B* originate from different overlapping binding domains in those regions.

Conversely, in *DupL* the duplicated region spans two TADs (**Fig. 2**). In our model, those TADs are produced by different prevailing types of binding sites. Accordingly, the portion of the duplication within the *Sox9* TAD (region B1’ in **Fig. 2C**) has enriched contacts with itself and its corresponding original sequence (B1), but less with the portion of the duplication within the flanking TAD and its original sequence (resp. region A2’ and A2). Since B1’ is enriched of self-contacts but has comparatively less interaction with its neighboring genomic regions A2’ and A2, it forms a separate chromatin domain (termed a ‘neo-TAD’^43^) remaining partially isolated from the rest, as seen in a snapshot of the 3D structure of the locus (**Fig. 2E**, red region). As the isolated neo-TAD does not include main genes, *DupL* has no phenotype^43^.

Finally, *DupC* produces a neo-TAD as much as *DupL*, however, it now includes a copy of the next flanking gene, *Kcnj2* (**Fig. S4**). As seen in the contact matrix of the full genome, within the neo-TAD the duplicated *Kcnj2* establishes ectopic contacts with the duplicated part of the regulatory region of *Sox9*. So, *Kcnj2* is mis-expressed, leading to the associated phenotype^43^.

In brief, our findings clarify how mutations impact chromatin architecture and the mode of action of the 3D structure in regulating gene activity. In particular, they explain how the considered structural genomic variations at the *Sox9* locus differently alter 3D conformation and gene regulation by specific enhancer hijackings, resulting in distinct phenotypes.

### Epigenetic profile of the binding domains

To shed light on the nature of the model inferred binding sites (**Fig. 1D**), we correlated their genomic locations with histone mark tracks available in the ENCODE database^49^for the GM12878 cell line. In particular, we used the binding domains derived from even-numbered chromosomes to compute such correlations, in order to use the derived barcode linking binding site types and epigenetics to later independently predict the architecture of odd-numbered chromosomes. In our analysis, we retained only statistically significant correlation values, i.e., those above a random control model with sites having bootstrapped genomic positions (Materials and Methods). As the different binding domains tend to fall in groups with similar epigenetic profiles, we clustered them to identify genome-wide significantly distinct epigenetic classes (Materials and Methods). The Akaike Information Criterion^50^ (AIC) returns a set of 9 statistically different groups (**Fig. 3A**), a result also supported by basic hierarchical clustering (**Fig. S6B**).

Three classes of binding domains strongly correlate with active chromatin marks (**Fig. 3A**), but they are distinct from an epigenetic point of view. While class 1 is enriched for only active marks, classes 2 and 3 are both enriched also in H3K9me3. Also, class 3 shows a stronger correlation with H3K4me1 as compared with class 2, a histone mark associated especially with active enhancer regions^10,44–47^. Interestingly, the genomic positions of the sites of the first three classes (**Fig. 3B**) are partially correlated (**Fig. S7C**, Materials and Methods). Their histone signatures are also consistent with DNA accessibility, early replication time and RNAseq transcription data (**Fig. S6C**). That supports the view that the binding sites in class 1, 2 and 3 are responsible, genome-wide, especially for specific contacts between transcribed and regulatory regions, mediated by factors such as active Pol-II, as experimentally demonstrated at a number of loci^40^. Class 4 has the typical signature of bivalent chromatin, with H3K27me3 combined with active marks. Its binding sites could be responsible for interactions between regions including, for instance, poised genes and their regulators, as seen in FISH co-localization experiments^40^. Classes 5 and 6 are significantly correlated with H3K27me3 and could be responsible for the experimentally observed self-interacting domains of PRC repressed chromatin^51^. Interestingly, the first six classes are the only ones to correlate with CTCF binding sites (**Fig. S6C**). That confirms the significance of CTCF in regulating chromatin architecture and gene activity (see, e.g. ^52^), highlighting that its role can be modulated by different sets of histone marks and molecular factors.

Classes 7 and 8 display a lack of active marks, but while class 8 does not correlate with any of the used histone marks, class 7 shows a correlation with H3K9me3, a mark usually associated with constitutive heterochromatin and lack of transcription factor binding. Finally, class 9 (named ‘low signal’) has a very low correlation with available histone marks. However, consistently with previous studies^10,44–47^, it covers almost 15% of the genome, while the other classes range from around 2% to 10% in genomic coverage (**Fig. S7A**). Interestingly, the different classes are significantly differently enriched over the different chromosomes and not consistent with a uniform random genomic distribution (**Fig. S7B**, p-values<0.05, Materials and Methods).

To understand the relative importance of the different types of binding domains in shaping chromatin architecture, we conducted a set of *in-silico* experiments with mutant models where each class, one at the time, is erased. Specifically, from the wild-type chromosome models we removed the binding domains of a given class. Next, we computed the contact maps of the mutated model and measured across chromosomes the variation of the Pearson, r, and distance-corrected Pearson correlation coefficient, r’, between the mutated model and wild-type Hi-C contact map (Materials and Methods). The variation is found to be proportional to the genomic coverage of the different classes in both cases (**Fig. S7D**,**E**). That implicates that no binding class has a special role in holding the architecture of the genome in place. The linear relation whereby the removal of, say, 10% of binding sites genome-wide roughly results in a 10% reduction of r highlights the structural stability of the system: the removal of a small fraction of binding sites proportionally alter the structure but does not produce a sudden collapse of the architecture, as reported by recent experiments^53–57^.

Finally, as a control of the robustness of the association between binding site types and epigenetics, we applied the same approach to the mentioned mouse ES cells^14^, using the corresponding set of ENCODE histone modifications in mouse, and found an overall analogous classification (**Fig. S8**).

Summarizing, the inferred binding site types have each a specific epigenetic barcode falling in classes that match well those found by previous epigenetic genome segmentation studies^10,44–47^. However, our binding domains are inferred from only Hi-C data without prior knowledge of epigenetics. Hence, they bring together independent information on architecture and epigenetics. A crucial feature of the model binding domains to explain contact data is that the different types do overlap with each other along the genome at the resolution of the considered Hi-C data. Therefore, they naturally provide each DNA window with a distinctive set of binding site types. This is an important difference with 1D epigenetic segmentation classes: by definition, those have no genomic overlap, thus each DNA window is associated to only one of such classes. Epigenetic segmentations have been shown, though, to correlate with Hi-C contacts^28,32,46^.

### Epigenetic linear segmentation only partially captures chromatin folding

To deepen our comprehension of the interplay of chromosome epigenetics and folding, we investigated the architectural information content retained in 1D epigenetic segmentations of the genome and compared it with the more complex DNA barcoding given by the classes of our binding domains. As done in previous studies^10,44–47^, we segmented chromosomes in 9 epigenetic classes based only on ENCODE histone marks (**Fig. 4A**,**B**). For simplicity, we opted for 9 classes to match the number of different types of binding domains found above. Such a number of classes is comparable to those in previous segmentation studies, and our results are not affected by more complex choices of segmentation (until the scale of the single binding domain is reached). Next, we derived *in-silico* the contact maps predicted by a polymer model based only on such a 1D epigenetic segmentation. Specifically, we considered a polymer model where chromatin physical interactions only occur between homologous 1D-segmented epigenetic regions^28^. Interestingly, while the overall contact patterns from such a model visually resemble Hi-C patterns (for example, r=0.78 for chromosome 20), their distance-corrected Pearson correlation, r’, with Hi-C data is very low (for chromosome 20 r’=0.02, **Fig. 4A**,**C** and **Fig. S9**, Materials and Methods). Hence, the patterns derived from a polymer model constructed from 1D epigenetic segmentation is only partially better than one where Hi-C pair-wise interactions are replaced by the average value corresponding to that genomic separation. Conversely, an SBS model with 9 types of binding domains, based on epigenetics classes, genomically overlapping as discussed before, has r=0.89 and r’=0.43 for chromosome 20; and, as stated, the model with the full set of inferred binding domains has r=0.97 and r’=0.84.

To understand the partial failure of 1D epigenetic segmentation in explaining contact data (**Fig. 4B**,**C**), for each pair of genomic sites we identified the binding domain that mostly contributes to their pair-wise interaction within the full SBS model (**Fig. 4D**,**E**,**F**, Materials and Methods). For clarity, we focus on a case-study 20Mb-wide region on chromosome 20. Plaid-patterns are visible in its Hi-C contact map, as expected from A/B compartments (**Fig. 4A**); they are also visible in the matrix of the most contributing binding domains (**Fig. 4F**), where rich and fine substructures appear as well. Consider, for instance, the TAD associated to region *C* in **Fig. 4**. The interactions within that TAD are mainly related to binding domains in class 7 (magenta, **Fig. 4F**), which is indeed the most abundant within the genomic region where *C* is located (**Fig. 4E**). Its interactions with the upstream region *A* can be simply traced back to homotypic interactions within class 7 itself, which is also the most abundant in A. However, the flanking region *B*, in which class 6 (dark blue) is the 1^st^ most abundant, also interacts with *C* (**Fig. 4F**). That occurs because class 7 is the 2^nd^ most abundant in *B* (**Fig. 4E**) and because in C class 6 is, in turn, the 2^nd^ most abundant. Such an example illustrates that a linear epigenetic segmentation model with homotypic interactions fails to account for the complexity of the observed contact pattern because a homotypic interaction between *B* and *C* would only occur if the two regions belong to the same class. Analogously, the contacts between regions *A* and *B* originate from different overlapping binding domains included in those regions. A similar reasoning can be extended to the plaid-pattern of A/B compartments (which is a specific example of a two classes genome 1D segmentation) capturing the overall interactions between homologous active and repressed regions respectively^7,42^. Yet, a much more complex and finer structure of contacts exists (including interactions across A and B compartments). Indeed, it has been shown that polymer models based on a linear epigenetic classification of domains are forced to include heterotypic interactions to accurately explain Hi-C data^32^.

Overall, homotypic interactions between the domains of a coarse-grained linear epigenetic segmentation of the genome, such as compartment A/B, are not enough to explain the specificity of Hi-C patterns with high accuracy since a complexity of relevant heterotypic contacts exists between those regions. The origin of those heterotypic interactions is understood within our analysis showing that multiple binding domains are present in a genomic segment. Their genomic 1D combinatorial overlaps associate a distinctive interaction profile to each DNA segment, containing the information required to produce through physics the complex details of the system 3D conformations (**Fig. 4**). In turn, the specific set of histone marks barcoding each binding domain provides a code linking epigenetic to architecture.

### The epigenetic barcode of binding domains predicts *de novo* chromatin architecture

To validate the identified association between linear epigenetic features and chromosome conformations, we considered a reverse approach whereby starting from only epigenetics data, through the mentioned barcode we identify the key binding sites of a set of independent chromosomes and, next, predict their contact matrices via polymer physics (**Fig. 5A**). Specifically, we exploited the epigenetic barcoding provided by the classification of the binding domains of even-numbered chromosomes, as previously described, to identify *de novo* the binding sites of odd-numbered chromosomes. To determine the locations and types of the binding sites, we partitioned each 5kb genomic window (5kb is the resolution of Hi-C) of odd-numbered chromosomes in equal-sized, 0.5kb sub-windows, which we epigenetically profiled by measuring the abundance of the mentioned key set of histone marks (Materials and Methods). We then computed the correlations between the epigenetic profile of each sub-window and the centroids of the epigenetic classes of the binding domains of even-numbered chromosomes (**Fig. 3A**). We focus on those epigenetics classes because they recapitulate the main functional groups found in segmentation studies; additionally, considering 9 types of sites is more stringent than considering all the binding domains found on even chromosomes. Exploiting such a larger set of domains would, of course, improve our results. Finally, each sub-windows of odd-numbered chromosomes was assigned with a binding site type corresponding to the epigenetic class having the highest correlation (**Fig. 5A**).

**Fig.5.**
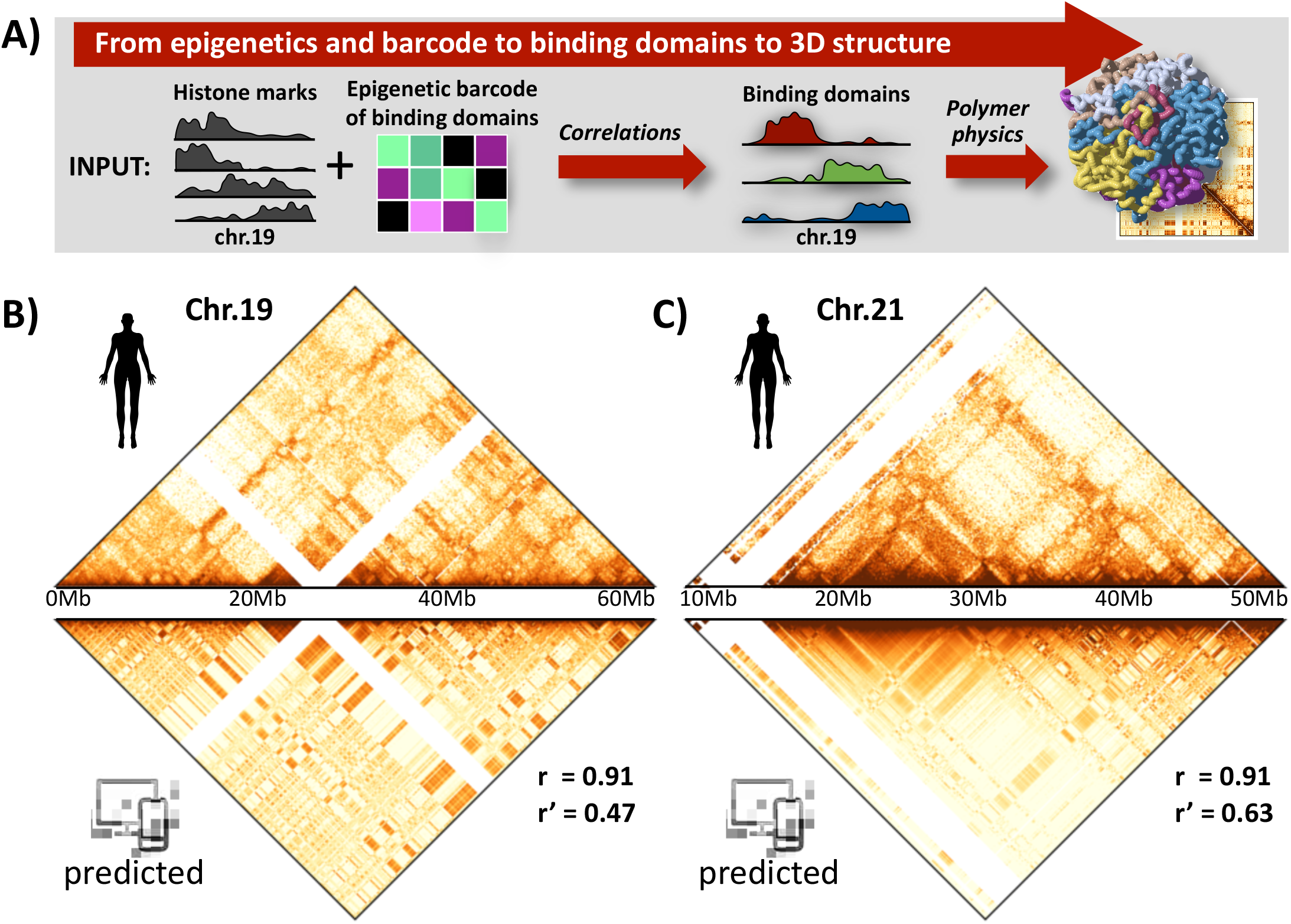
The epigenetic barcode of binding domains predicts chromatin contacts. **(A)** In a reverse approach, we correlate the epigenetic profiles of binding domains from even chromosomes with epigenetic signals from odd chromosomes to identify the binding sites of the latter. Next we use the SBS polymer model to predict 3D structures and contact matrices of odd chromosomes to be compared against independent Hi-C data. **(B)** Top: *in situ* Hi-C data^42^ (scales as in Fig. 1) of chromosome 19 in GM12878. Bottom: the predicted contact matrix has a correlation, a distance-corrected correlation and a stratum adjusted correlation with Hi-C respectively equal to r=0.91, r’=0.47 and SCC=0.65. **(C)** Top: Hi-C data of chromosome 21. Bottom: the predicted contact matrix has correlations with Hi-C equal to r=0.91, r’=0.63 and SCC=0.50.

Once obtained the genomic locations of the binding sites along odd-numbered chromosomes, we computed their contact matrices via the SBS polymer model and compared them with the corresponding *in situ* Hi-C maps (**Fig. S10A**,**B**). **Fig. 5B**,**C** shows, for example, the contact data of chromosomes 19 and 21 predicted by use of the above defined code that links binding sites, i.e., architecture, to epigenetic marks. In all the considered cases, the predicted matrices well capture the patterns of interactions seen in Hi-C data even at large genomic distances, albeit for simplicity we considered only 9 types of binding domains. Accordingly, the correlation and distance-corrected correlation coefficients (r=0.91 r’=0.47 and r=0.91 r’=0.63 for respectively chromosome 19 and 20) are much higher than those found by 1D epigenetic segmentation, as seen above.

Taken together, our results show that the barcode linking epigenetics marks to the binding domains inferred by PRISMR from Hi-C data, albeit still incomplete, can predict the genome’s 3D architecture to a good level of accuracy. A crucial difference between ours and epigenetic segmentation strategies to predict chromatin contacts^58^ is the intrinsically overlapping nature of binding domains, lacking in segmentations, which is necessary to recapitulate accurately the complex pattern of chromatin interactions.

## DISCUSSION

To infer from Hi-C data the different types of DNA binding sites determining chromosome architecture and their genomic position, we employed an approach based on machine learning^41^and the physics of the SBS polymer model of chromatin. The SBS model quantifies the scenario where TFs mediate the interactions between distal cognate binding sites, establishing DNA contacts and loops^21^. We found that the 3D structures derived by the model informed with the inferred putative binding domains, and folded through only polymer physics, explain Hi-C data genome-wide with high accuracy in human GM12878 B-lymphoblastoid^42^ and mES^14^ cells. That shows that the basic molecular ingredients considered by the model are sufficient to explain contact patterns across genomic scales. Thus, the binding domains encode key molecular information required to fold chromatin and provide an *architectural code* whereby 3D conformations can be established based on the 1D sequence (**Fig. 6**). To explain folding with high accuracy, they have a combinatorial organization along chromosomes, which is needed to control the intricate multitude of genomic interactions captured in Hi-C maps and their functional specificity, via a comparatively smaller number of molecular factors. Additionally, the non-trivial arrangement of binding domains provides structural stability to the 3D conformation of the genome, as experimentally reported^53–57^. We found that binding domains produce chromatin interactions extending across chromosomal scales, above the size of single TADs and A/B compartments, in a hierarchy of higher-order 3D structures, as in the meta-TADs^16^ picture.

**Fig.6.**
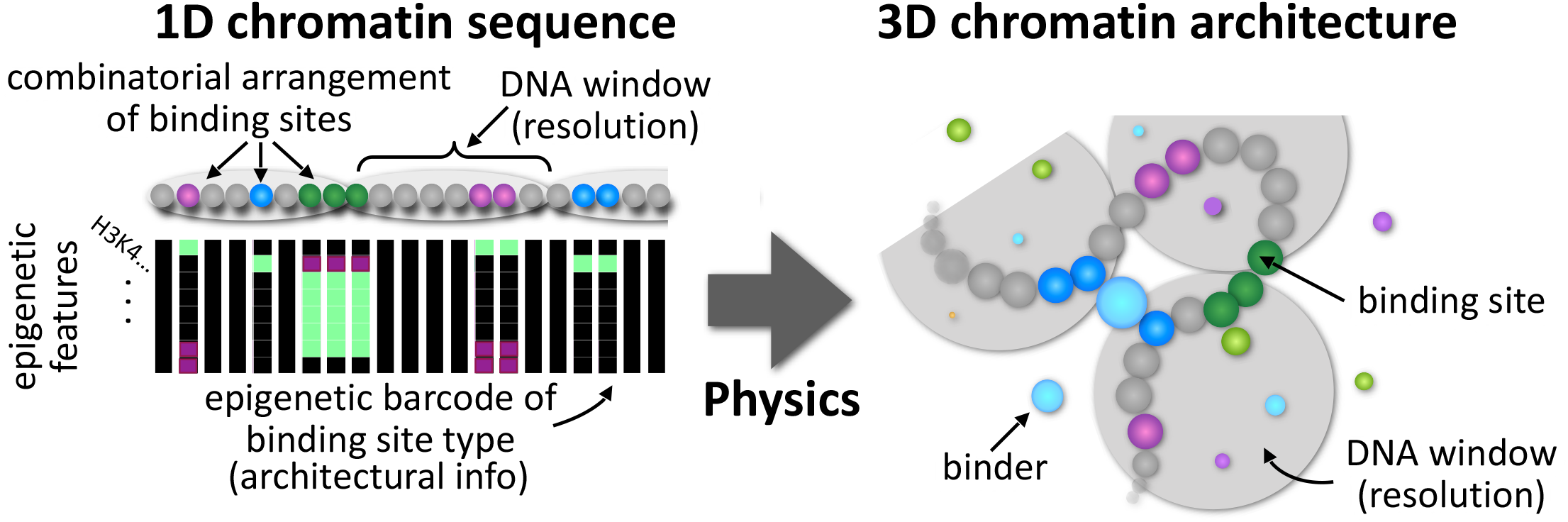
Chromatin 3D architectural information is encrypted in a combinatorial 1D arrangement of epigenetic barcoded sites. Our approach infers, from Hi-C data only, the minimal set of binding sites along the 1D genome sequence (left) required to produce, via polymer physics (e.g., interactions with diffusing cognate binding molecules), 3D structures (right) consistent with Hi-C contacts. The inferred binding sites are barcoded by specific epigenetic marks (vertical bars) and fall into epigenetic classes (bead color) well matching functional chromatin states known from linear segmentation studies. However, they have a genomic overlapping, combinatorial organization, lacking in epigenetic segmentations, necessary to explain Hi-C contacts with high accuracy genome-wide. Their epigenetic barcode was shown to predict *de novo* chromatin conformations, e.g., after genetic or epigenetic variations, showing that the inferred combinatorial 1D arrangement of binding sites carry accurate, specific 3D architectural information genome-wide.

Next, we associated each of the Hi-C inferred binding domains to an epigenetic profile based on the genomic correlation with a few main ENCODE histone marks. The model binding domains turn out to belong to main epigenetic classes, similar in human and mouse cell types, which well match known chromatin states (e.g., active, poised, repressed) derived by linear segmentation studies^10,44–47^. However, as stated, the identified binding domains have broad overlaps along the genome, a feature that is missing in linear segmentations but is required to explain Hi-C accurately. The few coarse-grained epigenetic classes here discussed constitute only a first, simplified description of the epigenetic features of the binding domains that shape chromatin architecture inferred by PRISMR. More generally, their barcode is expected to be associated with a broader set of (still partially unknown) molecular factors, including histone marks, CTCF^42^, Active/Poised Pol-II^40^ and additional factors, such as PRC1^51^, PRC2^40^and MLL3/4^59^. Furthermore, molecular mechanisms beyond those envisaged by the SBS model, such as DNA loop extrusion^22–24^, appear to play a role in chromosome folding and the code can be extended to accommodate them.

The inferred binding domains and the associated architectural interaction code were tested by making predictions on the changes of the 3D structure caused by a set of structural variants at the *Sox9* locus linked to human diseases. Notably, the predicted contact maps were confirmed by independent cHi-C data in cells carrying such mutations^43^. This is a stringent validation because there are no available fitting parameters. The model also helps understanding how the mutations differently affect the 3D structure of the locus (e.g., forming neo-TADs) and how that differently impacts gene regulation and, hence, phenotype by enhancer hijackings.

Finally, in a reverse approach, based on the discovered link between epigenetics and binding domains, we identified the binding sites of an independent set of chromosomes from only their epigenetic marks. Those binding sites were sufficient to predict *de novo*, via the physics of the SBS model, the contact matrices of those chromosomes with good accuracy, validating our approach.

Overall, the agreement between our results and independent experimental Hi-C data strengthens the scenario where chromatin 3D architectural information is encoded in a 1D combinatorial arrangement of epigenetically barcoded sites, which can be inferred across chromosomes and cell types by our computational approach. By integration of different genomic data, it provides a quantitative picture of the deep cause-effect relationship between epigenetics, architecture and function, which we tested to predict the phenotypic effect of mutations linked to congenital disorders. That can help the development of computational tools in biomedicine to infer the link between genotype and phenotype from the features of the genomic landscape.

## MATERIALS AND METHODS

### The String & Binders Switch model of chromatin

To investigate the 3D structure of the genome, we employed the String & Binders Switch (SBS) model^21,26,30^. According to the SBS, a chromatin filament (from small loci to entire chromosomes) is modeled as a self-avoiding walk polymer chain of beads, a fraction of which, named binding sites, interacts with diffusing molecular binders. The interaction between binding sites and binders allows for the formations of loops along the polymer and, therefore, permits its spontaneous folding (**Fig. 1C**). Each bead can be bound only by its specific, cognate type of binders and, to fully describe the complexity of the system, different types of interactions are allowed together with inert sites along the chain that do not interact with any binder (apart from steric effects). We represent these different interactions as different “colors” of the system, “gray” beads being the non-interacting particles (**Fig. 1C**). Key parameters of the model are the concentration, c, and the binding energy, **E**_int_, of each different type of binder. As a function of c and **E**_int_, the system of corresponding, cognate binding sites exhibits a coil-globule phase transition from an open conformation (at low concentration or energy) to a globule, compact phase (at high concentration or energy) as extensively discussed in previous studies^26,30,39^. The presence of different sets of binding sites (here named “binding domains” and represented with different colors) interacting with different, cognate molecular factors allows the formation of complex 3D structures by microphase separation.

### The PRISMR method

To determine the distribution of the different binding sites along the SBS polymer chain, here we used PRISMR, a previously illustrated machine learning procedure^41^. The PRISMR algorithm is a polymer physics-based method that, starting from an experimental contact matrix (e.g. Hi-C or GAM), finds the minimal polymer model that, at equilibrium, best describes the input. Although we focus on the SBS polymer model to describe a chromatin filament, the PRISMR algorithm can be easily generalized to different models.

A detailed description of the PRISMR method can be found in ref ^41^. Here we just summarize the key points of the algorithm. An SBS polymer model of a genomic region is composed of *L* beads, depending on the resolution of the input contact matrix of the region. For instance, a 10Mb locus at 10kb resolution is partitioned in *L*=1000 bins. Furthermore, we split each of the *L* bins into *r* different sub-units, considering that a single DNA bin could include many binding sites and interact with different factors. The SBS polymer is then completely characterized by the arrangement of the binding sites along the chain. Given the number n of different types of binding sites, PRISMR finds the color arrangement along the polymer chain by the minimization, via an iterative Simulated Annealing (SA) Monte Carlo optimization procedure^60,61^, of a specific cost function made of two terms. The first term representing the distance between the experimental and the model predicted contact matrices; the second one is a Bayesian term proportional to the total number of colored sites of the polymer through a parameter *λ* and penalizes the addition of new colored beads. In this way we account for the necessity to fit well the input data and, at the same time, we attempt to avoid overfitting. After initializing the SBS polymer in a random configuration, by assigning a random color to each bead, a standard iterative SA procedure is performed, as available in public software repositories (see e.g.^62^), to optimize the model^60,61^. Schematically, each SA step consists in randomly changing the color of a polymer bead, compute the average contact matrix of the new polymer, evaluate the new cost function, compare it with the cost function in the previous step and, based on it, accept or reject the color change. SA steps are iteratively repeated until convergence^41^. The entire procedure is repeated many times by varying the polymer initial configurations and the model parameters *n, r*, and *λ*, to find their optimal values.

### Details on the application of PRISMR genome-wide

In this study, we present the first genome-wide application of the algorithm. Precisely, here we applied PRISMR over the somatic chromosomes of the human genome, obtaining, for each chromosome independently, the SBS polymer that best describes its corresponding Hi-C matrix. We employed published in situ Hi-C data^42^ relative to the human GM12878 cell line at 5kb of resolution and normalized according to the method described in ref ^63^. To reduce the local noise in the input Hi-C data, we applied a gaussian filter with a standard deviation equal to 1 along both *x* and *y* directions. The optimal value of the parameters of the algorithm has been estimated as already described in ref ^41^, that is, we repeated the SA procedure many times starting from different initial conditions and different values of *n, r*, and *λ* to set these parameters at the values that explain the input data within a given accuracy. As input data for the optimal parameter evaluation, we used the contact matrix of chromosome 12, a medium-sized chromosome, obtaining *n*=30 different types of binding sites, *r*=30 and *λ*=3×10-5. The same values for the parameters *n, r*, and *λ* have been used to obtain the best SBS polymer for all the other chromosomes. **Fig. S1A** shows the comparison between the contact matrices inferred by PRISMR (lower triangular maps) and the in situ Hi-C matrices (upper triangular maps). The global pattern obtained by PRISMR is highly correlated with the experimental one as also quantified by the comparatively high values of the Pearson’s (r), distance-corrected Pearson’s (r’)^41^ and stratum-adjusted (SCC)^48^ correlation coefficients (**Fig. S1B**, see below). In the calculation of r and r’, to correct for outliers, we did not consider genomic distances below 25kb. The PRISMR method is highly generalizable across different experiments and data resolution. To test that, we also applied our method to genome-wide Hi-C data in mouse embryonic stem (mES) cells^14^ at 40kb resolution (**Fig. S2A**). The correlations between experimental and model matrices obtained in mouse are as high as the values obtained in human, as shown in **Fig. S2B**.

### Structural variants at the Sox9 locus and validation of PRISMR

As a validation of the PRISMR inference method and the SBS model, we implemented in-silico a set of three previously studied structural variants in E12.5 limb bud cells from mice^43^. Specifically, we started from a SBS polymer model3 of the region chr11:109000000-115000000 (mm9, mESC cells) including the *Sox9* gene and implemented on it, independently, the following duplications: *DupS*, an intra-TAD duplication of the region chr11:111760000-112160000; *DupL*, an inter-TAD duplication of the region chr11:110960000-112520000; *DupC*, another inter-TAD duplication of the region chr11:110760000-112520000. We then computed the PRISMR predicted contact maps for each duplication, under no adjustable parameters, obtaining the following values of correlations r and r’, between model and experimental matrices (excluding the effect of strong outliers <5th and >95th percentile): r=0.88 and r’=0.48 in *DupS*; r=0.82 and r’=0.41 in *DupL*; r=0.82 and r’=0.47 in *DupC* (**Fig. S4**).

### Matrix similarity evaluation

The agreement between experiment and model matrices has been quantified using Pearson’s correlation coefficient, r. We also used two additional measures: 1) the distance corrected Pearson correlation coefficient, denoted by r’, that is the Pearson’s correlation coefficient between the two matrices where we subtracted from each diagonal (corresponding to a given genomic distance) their average contact frequency; 2) the stratum-adjusted correlation coefficient, denoted by SCC, from the HiCRep^48^ method with a smoothing parameter h=10 and an upper bound of interaction distance equal to 5Mb. These two measures have been used to put aside the expected decreasing trend of the pairwise contact frequency with genomic distance, which tends to dominate in the simple Pearson correlation value.

### Molecular Dynamics simulations

To obtain 3D conformations of the PRISMR derived SBS models, shown in **Fig. 1F, Fig. 2D,E** and **Fig. S5C,D**, we performed Molecular Dynamics (MD) simulations. To this aim, we proceeded as described in ref ^30^. Briefly, the polymer chain and the binders move in the system according to the Langevin equation, integrated with the LAMMPS software^64^, using standard dimensionless parameters^65^. The SBS parameters used are the same reported in ref ^30^, i.e., the beads and binders interact with an interaction energy **E**_Int_=8.1KbT and the binder concentration is high enough to allow the coil-globule transition (c=194nmol/l for the *Sox9* WT and similar values for the duplications). To make MD computation times feasible for the entire chromosome 20, we considered a coarse-grained version of its SBS polymer, having a 50-fold reduced number of beads. All the conformations are taken in the equilibrium globular phase. In all the snapshots, beads coordinates have been interpolated with a smooth third-order polynomial splice curve by using the POV-RAY^66^ software.

### Characterization of the binding domains arrangement along chromosomes

To study how the different binding domains (colors) span along the genome, we employed two main measures. The first one, that measures the domain size, is the genomic coverage, i.e., the fraction of beads of a given color multiplied by the length of the chromosome it belongs to. Averaging over all the sizes of the domains identified by PRIMSR across chromosomes, we find that the genomic length covered by each domain is on average 3.1 Mb, with a standard deviation of 1.9 Mb, a value close to the mean-size of a TAD. To measure, instead, the range of the interactions due to a single binding domain, we defined *r*_*Int*_ as two times the standard deviation of the center of mass of that domain. The distribution of *r*_*Int*_, P(*r*_*Int*_), extends far beyond the size of the single domain, ranging from a few mega-bases to more than 100 Mb (**Fig. S3A**). To check the statistical significance of the domains identified by PRISMR, we compared P(*r*_*Int*_) with a control model obtained by randomly bootstrapping the location of our binding sites along the genome, and we found that the two distributions are significantly different (p-value<0.001, Wilcoxon’s rank sum test). We also found that P(*r*_*Int*_) is asymptotically consistent with a power-law scaling, as shown in **Fig. S3A** where the right-hand side of the distribution is well described by a power-law fit (dotted red curve in the graph).

Another way to test the significance of the binding domains identified by PRISMR is to measure their mutual overlap^41^, to be compared with the expected level of overlap in the random model of bootstrapped domains mentioned before. To this aim, given a pair of different domains on a chromosome, we defined their overlap *q* as the sum of products of binding sites occurrences of the two colors in each genomic window, normalized to have *q*=100% in the case of identical domains (the cartoon in **Fig. S3B** gives a visual impression of what *q* is measuring). We found that the distribution P(*q*) of the overlap of the binding domains predicted by PRISMR is significantly different (p-value<0.001, Wilcoxon’s rank sum test) from the one expected in the random control model (red and blue distributions in **Fig. S3B**, respectively).

### Epigenetic analysis of the binding domains

To obtain insight into their molecular nature, we analyzed the PRISMR inferred binding domains in the light of epigenetics data. To this aim, we downloaded from the ENCODE database^49^ a set of 5 key histone modifications (H3K4me3, H3K4me1, H3K36me3, H3K9me3 and H3K27me3) in the human GM12878 cell line. ChIP-Seq signals were binned at 5kb resolution by summing the signal contained within each 5kb window (using the bedtools map tool from the bedtools^67^ software). After that, to measure the similarity between our binding domains and the histone marks, we computed Pearson’s correlation coefficient between the number of binding sites of each domain and each histone mark profile. Next, we employed a control model to retain only statistically significant correlations. To this aim, first, we computed the Pearson correlations between chromatin mark signals and randomized binding domains signals obtained by bootstrapping their actual genomic locations; then, we retained as significant only the correlation values above the 95th or below the 5th percentile of the distribution of the random correlations. We then collected data in a rectangular matrix *X*, whose element *X*_*ij*_is either the significant correlation between the *i*-th binding domains and the *j*-th histone mark or zero if the correlation was not significant. Since each row of *X* represents a binding domain’s correlation profile with the considered histone modifications, we refer to them as the epigenomic signature of the binding domain. To find binding domains with similar epigenomic signatures, we performed a hierarchical clustering analysis on *X* using the *Python SciPy* clustering package with ‘Euclidean’ distance metric and ‘Ward’ linkage method. To assess the number of clusters in the hierarchical clustering output, we cut the dendrogram at different values (ranging from one to the number of binding domains) and evaluated the Akaike Information Criterion^50^(AIC) as the number of clusters *k* is varied. As shown in **Fig. S6A**, while no sharp transitions are present, the curve has a global minimum at *k*=9. We therefore grouped all the different rows of *X* in 9 different classes according to their affinity to each cluster (**Fig. S6B**). Each of the 9 classes can be characterized by the epigenetic signature of its centroid, which is the average histone signature of the domains belonging to the given class (**Fig. 3A**). To assign biologically meaningful labels to the obtained classification, we looked at the enrichment of several types of functional annotations. Precisely, we first binned each annotation track at 5kb resolution, then, for each pair of annotation mark and epigenetic class, we computed the average of the Pearson correlation values between that mark and the binding domains of that class (see **Fig. S6C**). The set of functional annotations in GM12878 cell line considered in this study is taken from ENCODE and include: (1) all remaining available histone modifications; (2) transcription factors binding sites; (3) DNase hypersensitive sites; (4) replication timing data from the Repli-seq assay; (5) transcription data from RNA-seq assay (**Fig. S6C**).

To further test the association between binding domains and epigenetics, we repeated the above analysis for the mouse case. Specifically, we computed correlations among the genome-wide binding domains obtained from Hi-C data in mES cells and a corresponding set of ENCODE histone modifications in that cell line. As shown in **Fig. S8A-C**, we found an overall similar epigenetic classification of the binding domains in human and mouse.

### Characterization of epigenetic classes of binding domains

The genomic coverage of a given epigenetic class has been computed as the fraction of sites of the binding domains belonging to that class (**Fig. S7A**). To study, instead, how the domains of a given class are distributed along the chromosomes, we counted, for each class, the number of domains falling in each chromosome (**Fig. S7B**, dotted lines are the average values). We found that their distribution is significantly different over the different chromosomes, as measured by the comparison with a uniform distribution obtained by randomly bootstrapping the domains of a given class over the chromosomes (p-value<0.05 for each epigenetic class, Kolmogorov-Smirnov test). We also asked whether the genomic positions of the sites of the different classes (**Fig. 3B**) were correlated with each other. To figure out that, we computed the Pearson correlation between the genomic location of the sites of all the possible pairs of epigenetic classes, averaged over the different chromosomes (**Fig. S7C**). We found that classes with similar histone signature correlate with each other and anti-correlate with classes showing a very different histone pattern.

We investigated the impact of the different epigenetic classes on genome architecture by measuring the effect on contact matrices of the withdrawal of the binding domains belonging to each class. Precisely, given the list of the binding domains of a class, we replaced their interacting binding sites with gray, non-interacting elements along each chromosome. We then computed the PRISMR contact matrices of the modified SBS polymer and measured their correlations r and r’ with Hi-C. Finally, we evaluated the variation of the correlation, Δr and Δr’, with respect to the wild-type model (r=0.94 and r’=0.76), averaged over all chromosomes. The variations of r and r’ obtained are shown as a function of the genomic coverage of each epigenetic class in **Fig. S7D**.

### Most abundant and most contributing binding domains to chromatin pairwise contacts

As the different binding domains can overlap with each other, to better visualize their locations along the genome, we show in **Fig. 4E** (upper bar) the 1^st^ most abundant binding domain, i.e. the one with the largest number of binding sites, per bin. Analogously, **Fig. 4E** (lower bar) shows the 2^nd^ most abundant binding domain per bin. In both cases, to help the visualization, the domains are colored with their epigenetic class color.

The contribution of the different binding domains in forming the interactions between bin pairs is then highlighted in **Fig. 4F**, where the colors of the most contributing binding domains are shown. Specifically, for a given pair-wise contact, we defined the contribution of a binding domain to that contact as the number of pairs of its binding site type between the two considered bins. The binding domain having the highest number of binding site pairs is the most contributing one and is colored with the color corresponding to its epigenetic class.

### Epigenetic linear segmentation model

To obtain a model based exclusively on the interaction among segments with a similar epigenetic profile, we considered the dataset of five histone modifications discussed in section “*Epigenetic analysis of the binding domains*”. We marked each 5kb genomic window with the z-score value of the signal of each histone mark in that window. Then, we performed a hierarchical clustering analysis to gather the genomic windows with similar histone profiles in 9 different groups, in order to match them with the 9 different types of binding domains found above. The obtained linear segmentation has been employed to define a polymer model for chr.20 with 9 different colors corresponding to the different linear epigenetic classes (**Fig. 4B**), where interactions can only occur between same-colored windows. Finally, we derived in-silico the contact map of such a model and compared it with the corresponding experimental matrix (**Fig. 4A-C** and **Fig. S9A-B**). We found that the Pearson correlation and distance-corrected Pearson correlation between the matrices are r=0.80 and r’=0.21.

We have also considered an additional model by assigning each of the different binding sites of chr.20 the color of the epigenetic class it belongs to. We found that this 9 color SBS model, that in contrast to the linear segmentation model has overlapping binding domains, has correlations r=0.89 and r’=0.43 with Hi-C.

### Prediction of *de novo* chromatin structures from epigenetic data by combinatorial barcode

The derived combinatorial code linking 3D conformation to 1D epigenetic signature can be used to predict de novo binding domains in independent chromosomes from epigenetics data only. Specifically, we used the code derived from the set of even-numbered chromosomes in GM12878 to predict the location of the binding sites along the odd-numbered chromosomes in the same cell line. To this aim, we partitioned each of their 5kb windows (which is the in situ Hi-C data resolution) in ten 500-bp sub-windows and binned the signal of the five key histone marks (H3K4me3, H3K4me1, H3K36me3, H3K9me3 and H3K27me3) in those sub-windows. In this way, we obtained a state vector for each sub-window, whose components are the histone marks’ abundances in that window. We checked that different sub-windows partitions, ranging from 5 to 20 sub-windows per bin, led to only marginally different results. To assign each sub-window to a specific binding site type (in the sense of the SBS model), we compared them with the centroids of the epigenetic classes of the binding domains of even-numbered chromosomes. Precisely, we computed the Pearson correlation coefficient between the state vector and each row of the centroid matrix, then assigned to that sub-window a binding site type corresponding to the epigenetic class with the highest correlation. Besides, two non-interacting ‘gray’ beads were added in each sub-window, so to match the number of beads per 5kb-bin of the PRISMR inferred polymer models. The described procedure results in an SBS polymer with 9 different binding domains for each odd chromosome. Afterward, we used the SBS model to calculate the predicted polymers’ contact matrices and compared them with the independent Hi-C data (Fig. **S10**). As reflected by the Pearson and distance corrected Pearson correlations, in all cases, the contact pattern is well described (see for instance chromosomes 19 and 21 in **Fig. 5**).

## ACKNOWLEDGMENTS

M.N. acknowledges support from the NIH grant ID 1U54DK107977-01 and 1UM1HG011585-01, the EU H2020 Marie Curie ITN n.813282, CINECA ISCRA ID HP10CYFPS5 and HP10CRTY8P, Einstein BIH Fellowship Award (EVF-BIH-2016 and 2019), Regione Campania SATIN Project 2018-2020. S.B. and A.M.C. acknowledge support from the CINECA ISCRA grant ID HP10CCZ4KN. We acknowledge computer resources from INFN, CINECA, ENEA CRESCO/ENEAGRID^68^and *Scope*/*ReCAS/Ibisco* at the University of Naples.

## AUTHOR CONTRIBUTIONS

MN designed the project. AE, SB developed modeling. AE, SB, AMC, AA, LF, MC, RC run the computer simulations and performed analyses. MN, AE, SB, AMC wrote the manuscript.

## COMPETING INTERESTS STATEMENT

The authors declare no competing interests.

## Notes

### Competing Interest Statement

The authors have declared no competing interest.

