## Supplementary Material for "Polymer physics and machine learning reveal a combinatorial code linking chromatin 3D architecture to 1D epigenetics"

A)

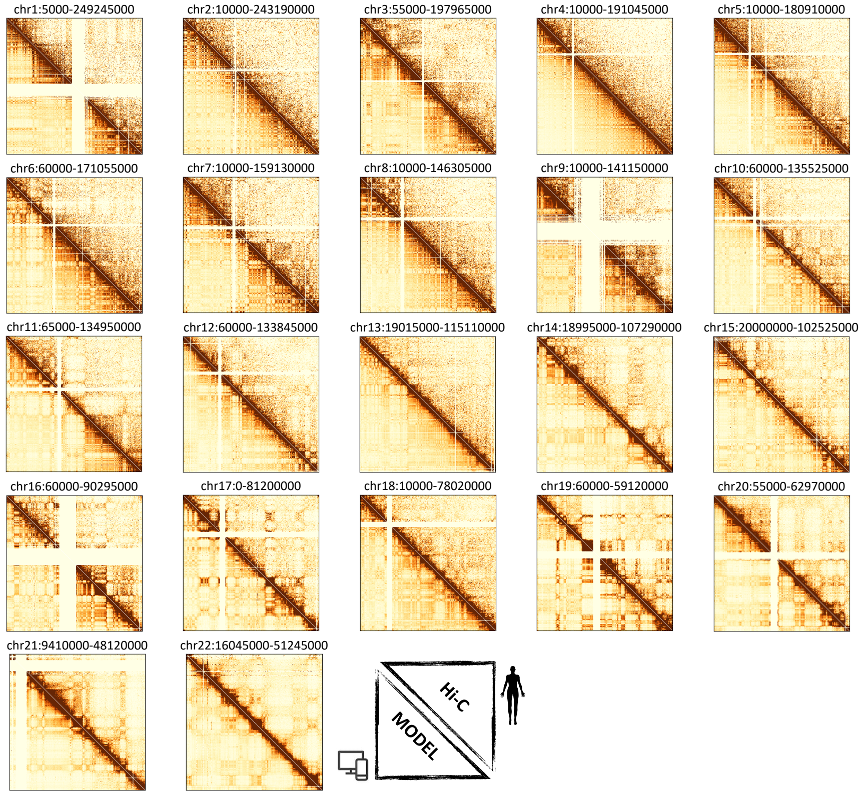


B)

| Chr | *r* | *r'* | SCC |
|---|---|---|---|
| 1 | 0.92 | 0.71 | 0.82 |
| 2 | 0.92 | 0.67 | 0.84 |
| 3 | 0.95 | 0.75 | 0.86 |
| 4 | 0.92 | 0.75 | 0.86 |
| 5 | 0.92 | 0.68 | 0.85 |
| 6 | 0.95 | 0.76 | 0.86 |
| 7 | 0.93 | 0.67 | 0.84 |
| 8 | 0.93 | 0.71 | 0.82 |
| 9 | 0.85 | 0.63 | 0.77 |
| 10 | 0.91 | 0.64 | 0.80 |
| 11 | 0.95 | 0.75 | 0.83 |
| 12 | 0.94 | 0.80 | 0.87 |
| 13 | 0.97 | 0.80 | 0.90 |
| 14 | 0.96 | 0.78 | 0.90 |
| 15 | 0.90 | 0.57 | 0.81 |
| 16 | 0.95 | 0.82 | 0.86 |
| 17 | 0.95 | 0.72 | 0.89 |
| 18 | 0.98 | 0.86 | 0.90 |
| 19 | 0.97 | 0.85 | 0.94 |
| 20 | 0.97 | 0.85 | 0.92 |
| 21 | 0.97 | 0.91 | 0.92 |
| 22 | 0.95 | 0.70 | 0.88 |

**Fig. S1**

**(A)** Contact maps (scales as in Fig. 1) across chromosomes from the PRISMR inferred SBS model (lower triangle) and from Rao et al. 2014 in situ Hi-C data in GM12878 (upper triangle). **(B)** Pearson, distance-corrected Pearson and stratum adjusted (SCC) correlation coefficients (respectively) between model and in situ Hi-C data. SCC values were computed using HiCRep (Yang et al. 2017).

A)

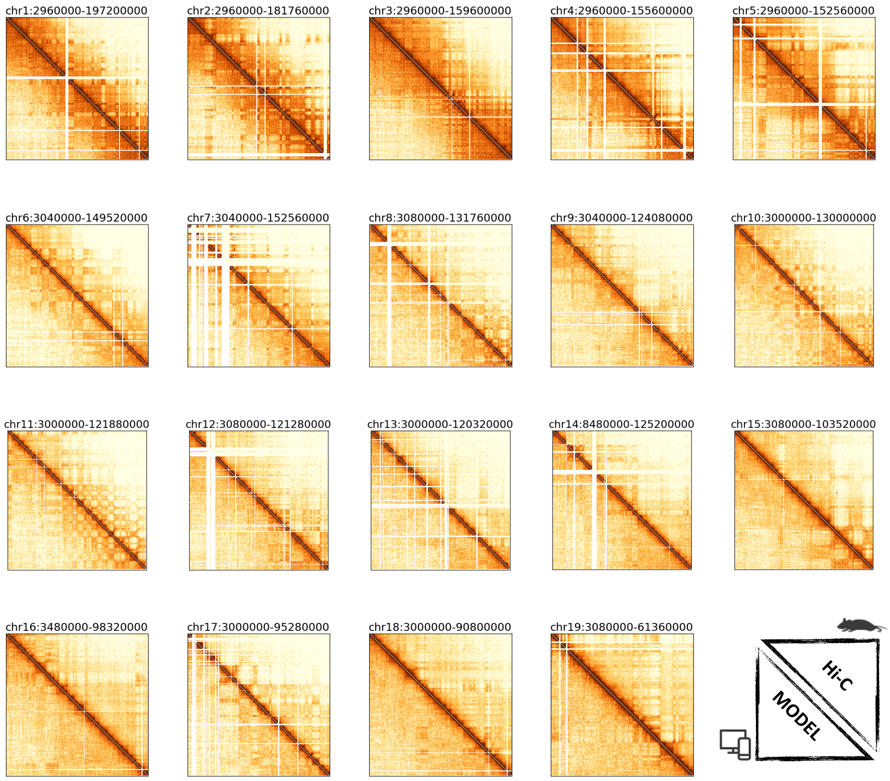


B)

| Chr. | *r* | *r′* | SCC |
|---|---|---|---|
| 1 | 0.94 | 0.55 | 0.85 |
| 2 | 0.94 | 0.59 | 0.86 |
| 3 | 0.94 | 0.59 | 0.84 |
| 4 | 0.95 | 0.64 | 0.89 |
| 5 | 0.94 | 0.62 | 0.87 |
| 6 | 0.98 | 0.73 | 0.89 |
| 7 | 0.96 | 0.68 | 0.88 |
| 8 | 0.94 | 0.55 | 0.81 |
| 9 | 0.94 | 0.55 | 0.78 |
| 10 | 0.94 | 0.52 | 0.77 |
| 11 | 0.95 | 0.60 | 0.83 |
| 12 | 0.94 | 0.58 | 0.81 |
| 13 | 0.94 | 0.54 | 0.77 |
| 14 | 0.95 | 0.56 | 0.79 |
| 15 | 0.95 | 0.58 | 0.77 |
| 16 | 0.94 | 0.55 | 0.76 |
| 17 | 0.95 | 0.61 | 0.82 |
| 18 | 0.95 | 0.58 | 0.77 |
| 19 | 0.96 | 0.69 | 0.81 |

**Fig. S2**

**(A)** Contact maps (scales as in Fig. 1) across chromosomes from the PRISMR inferred SBS model (lower triangle) and from Dixon et al. 2012 Hi-C data in mESC (upper triangle). **(B)** Pearson, distance-corrected Pearson and stratum adjusted (SCC) correlation coefficients (respectively) between model and in situ Hi-C data. SCC values were computed using HiCRep (Yang et al. 2017).

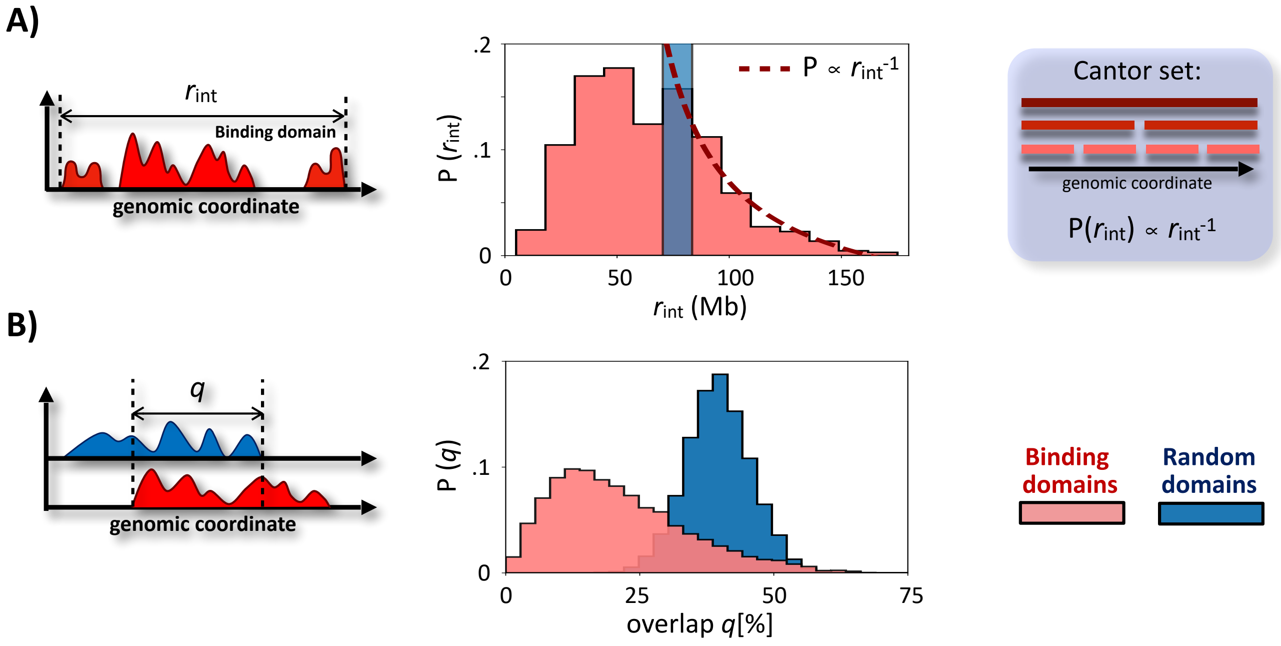


**Fig. S3**

**(A)** Distribution of the range of interaction of the PRISMR inferred SBS binding domains genome wide. The blue bar corresponds to a random model where the binding sites are bootstrapped. A Cantor set has hierarchically nested domains: the distribution of their ranges scales as an inverse power law. **(B)** The distribution of overlaps between the model binding domains compared to the one expected in the mentioned random model (blue).

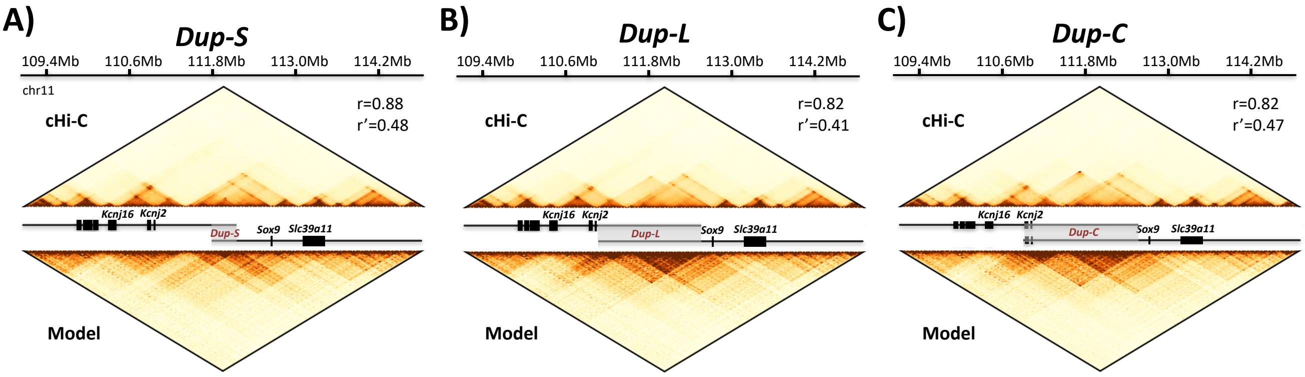


**Fig. S4**

**(A)-(C)** cHi-C data (Franke et al. 2016**,** top) and model predictions (bottom) across the available mutations in the *Sox9* locus, along with the Pearson, r, and distance-corrected Pearson, r’, correlation coefficients between model predictions and cHi-C data.

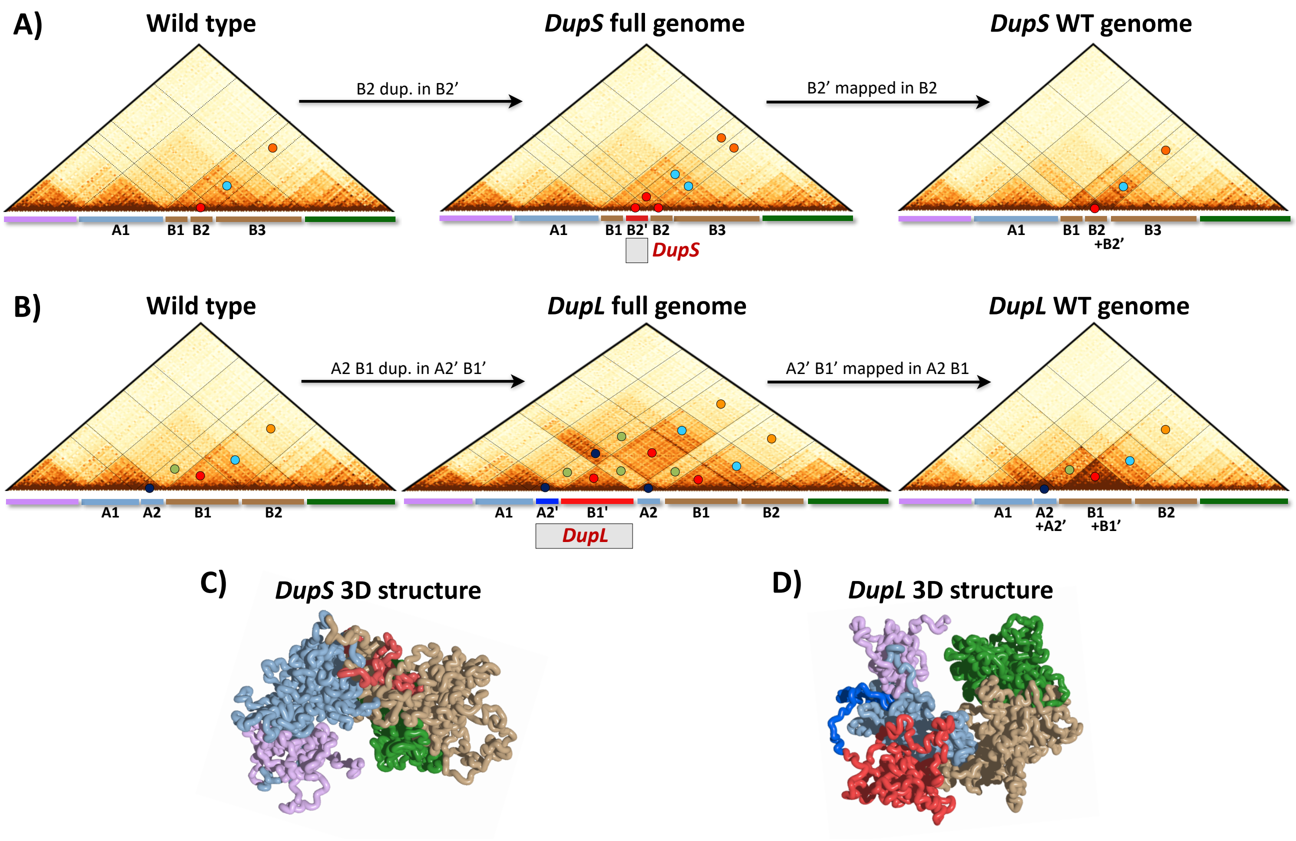


**Fig. S5**

Mapping the SBS model predicted contacts on the **(A)** *DupS* and **(B)** *DupL* full genomes clarifies the origin of the novel interactions and of neo-TAD discovered in *DupL* (Franke et al. 2016). The colored circles help visualizing the different regions of interactions of the duplicated sequences and how they map onto the wild-type genome, as reported in Hi-C experiments. **(C)-(D)** Model predicted 3D conformation of the mutated loci. Panels B and D are also shown in Fig.2, reported here to help the comparison between the two mutations.

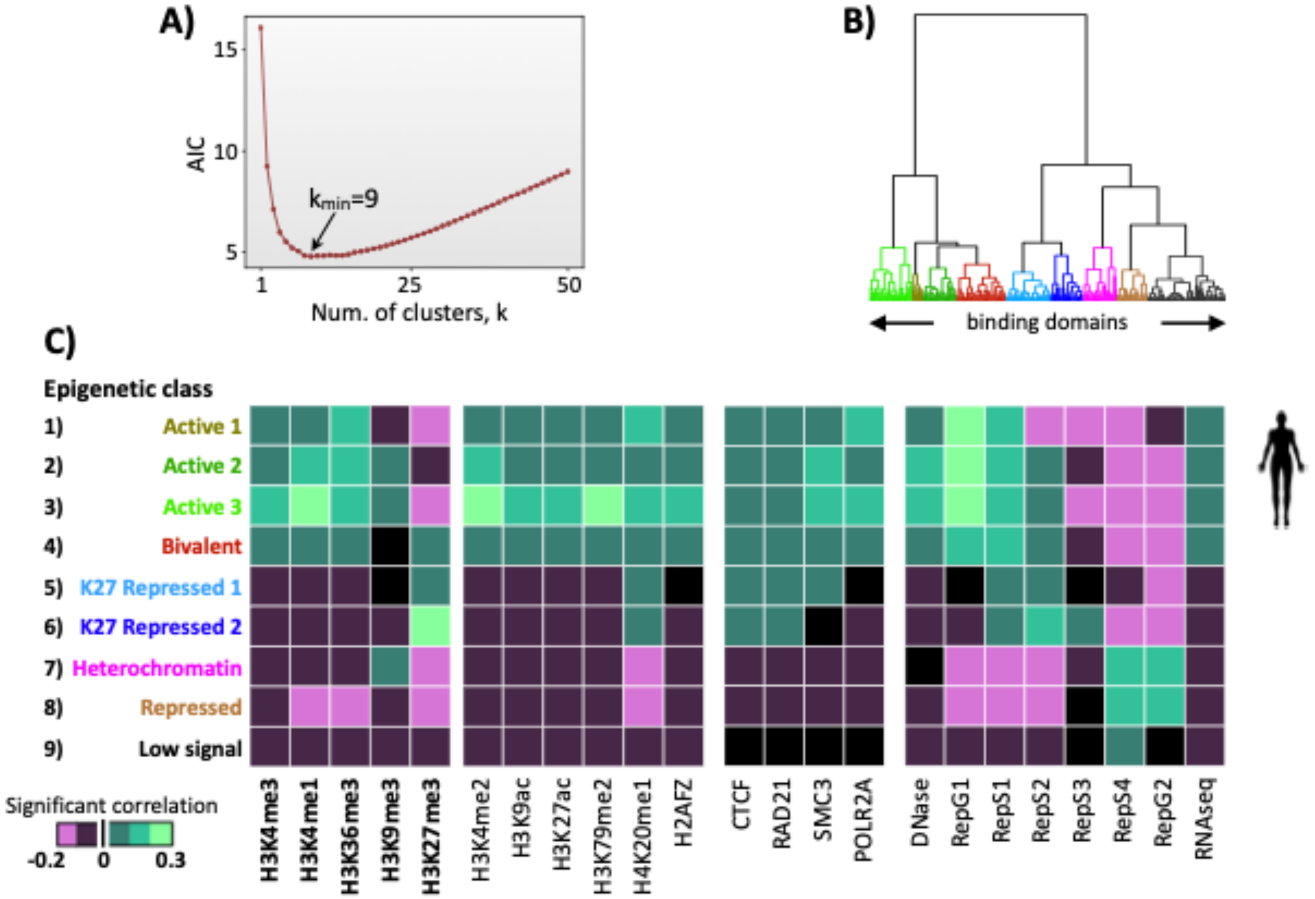


**Fig. S6**

Epigenetic classification of the binding domains in human GM12878 cell line: **(A)** The AIC statistical criterion has a minimum at k=9 clusters of binding domains based on their epigenetic profile shown in panel C. **(B)** A hierarchical clustering of the PRISMR inferred SBS model binding domains with the 9 identified classes highlighted. **(C)** The epigenetic signature of the 9 classes and their significant correlations with histone modification, transcription factors, DNA accessibility, DNA replication time and expression data.

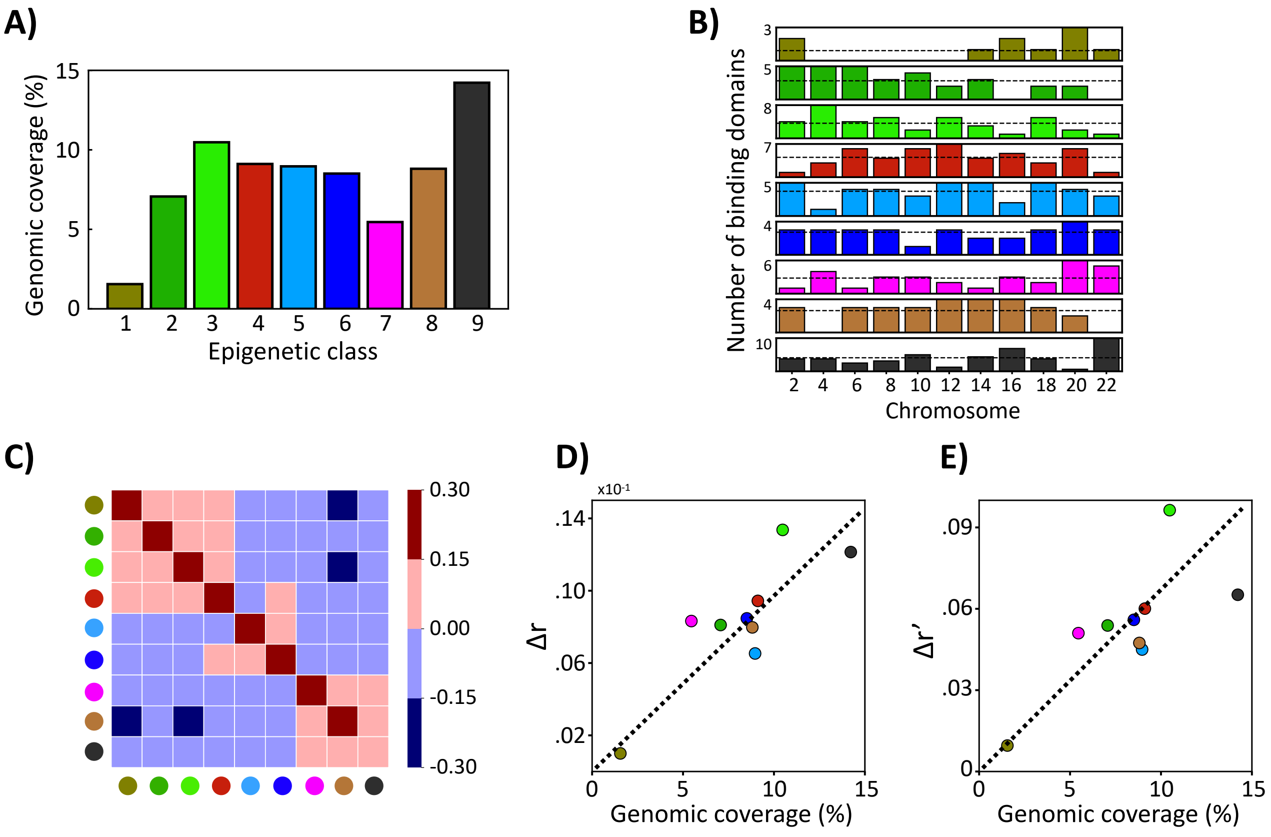


**Fig. S7**

**(A)** Genomic coverage of the 9 main epigenetic classes of the SBS model binding domains. **(B)** Relative number of the binding domains of the different classes across chromosomes. The distribution is not uniform (p<0.05). **(C)** Pearson correlation coefficient of the genomic location of the different classes over chromosomes. **(D)-(E)** Effect of the withdraw of a class of binding sites as a function of its genomic coverage. The effect of a class removal on the architecture is measured by the variation of the Pearson, r (panel D), and distance corrected Pearson, r’ (panel E), correlation with respect to the wild-type model. Δr’ is the difference between r’ in the wild-type model (r’=0.76) and in a model where the domains of a given class are removed, averaged over chromosomes. Analogously, Δr, is the wild-type r=0.94 minus r in the mutated model.

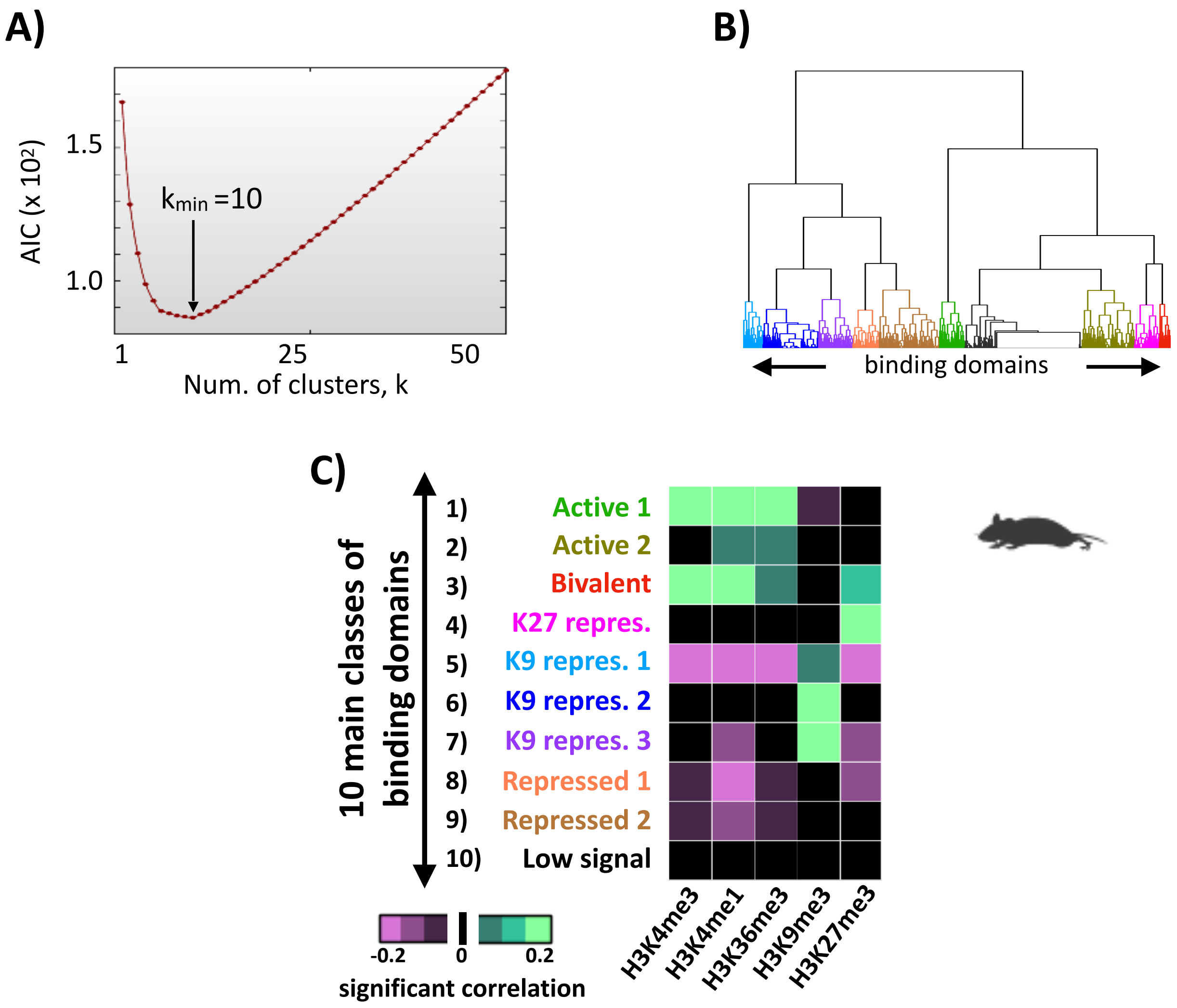


**Fig. S8**

Epigenetic classification of the binding domains in mouse embryonic stem cells: **(A)** The AIC statistical criterion has a minimum at k=10 clusters of binding domains based on their epigenetic profile shown in panel C. **(B)** A hierarchical clustering of the PRISMR inferred SBS model binding domains with the 10 identified classes highlighted. **(C)** The epigenetic signature of the 10 classes.

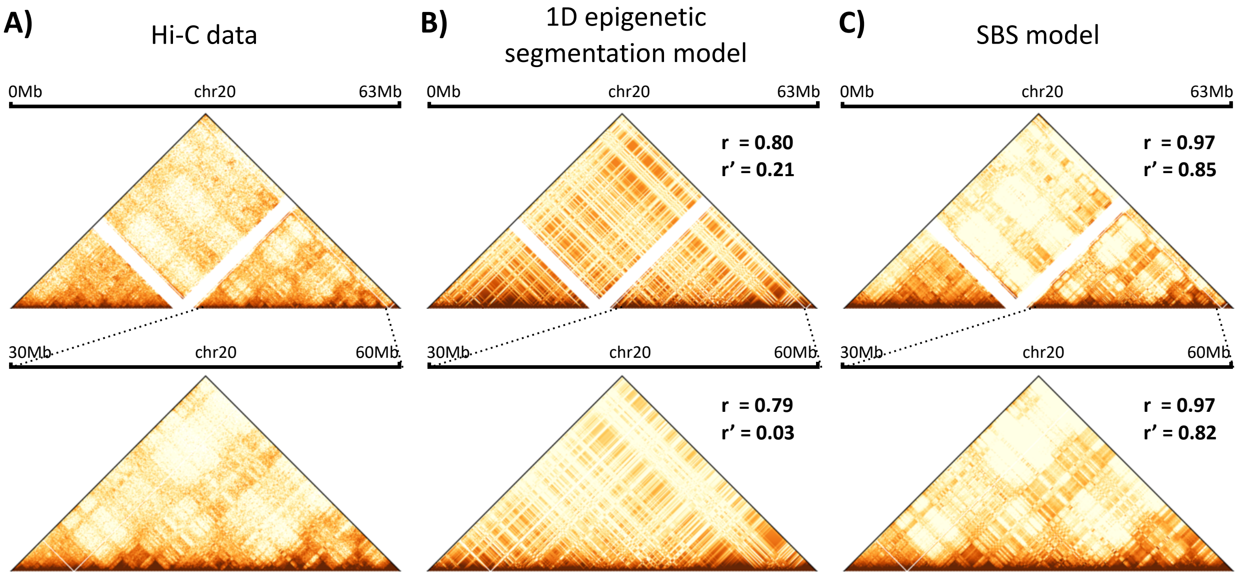


**Fig. S9**

**(A)** In situ Hi-C data in GM12878 (Rao et al. 2014) of the entire chr.20 (top) and of a zoomed 30Mb wide region (bottom). **(B)** Contact maps from a model of the chr.20 based only on homotypic interactions between linear segmented epigenetic regions. Its Pearson correlation, r, and distance corrected Pearson correlation, r', with the Hi-C matrices of panel A is shown for both regions. **(C)** The PRISMR inferred SBS model contact map for those regions together with the corresponding correlations.

A)

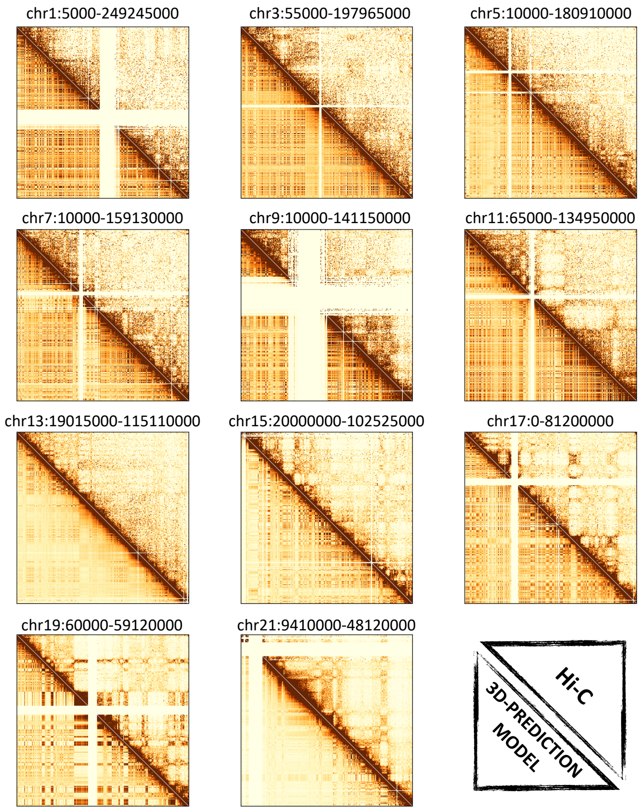


B)

| Chr | *r* | *r'* |
|---|---|---|
| 1 | 0.90 | 0.60 |
| 3 | 0.91 | 0.34 |
| 5 | 0.89 | 0.34 |
| 7 | 0.89 | 0.35 |
| 9 | 0.80 | 0.47 |
| 11 | 0.91 | 0.39 |
| 13 | 0.93 | 0.28 |
| 15 | 0.83 | 0.15 |
| 17 | 0.91 | 0.44 |
| 19 | 0.91 | 0.47 |
| 21 | 0.91 | 0.63 |

**Fig. S10**

**(A)** The upper triangles show the in situ Hi-C maps from the odd-numbered chromosomes in GM12878, while the lower triangles show the contact maps obtained by the predicted polymer models (scales as in Fig. 1). **(B)** Pearson and distance-corrected Pearson correlation coefficients (respectively) between the predicted matrices and the corresponding in situ Hi-C data.
